# *Plasmodium* male gametocyte development and transmission are critically regulated by general and transmission-specific members of the CAF1/CCR4/NOT complex

**DOI:** 10.1101/350116

**Authors:** Kevin J. Hart, Jenna Oberstaller, Michael P. Walker, Allen M. Minns, Mark F. Kennedy, Ian Padykula, John H. Adams, Scott E. Lindner

## Abstract

With relatively few known specific transcription factors to control the abundance of specific mRNAs, *Plasmodium* parasites also regulate the stability and turnover of transcripts to provide more comprehensive gene regulation. *Plasmodium* transmission stages impose translational repression on specific transcripts in part to accomplish this. However, few proteins are known to participate in this process, and those that are characterized primarily affect female gametocytes. We have identified and characterized PyCCR4-1, a putative deadenylase, which plays a role in the development and activation of male gametocytes, regulates the abundance of specific mRNAs in gametocytes, and ultimately increases the efficiency of host-to-vector transmission. We find that when *pyccr4-1* is deleted or its protein made catalytically inactive, there is a loss in the initial coordination of male gametocyte maturation and a reduction of parasite infectivity of the mosquito. Expression of only the N-terminal CAF1 domain of the essential CAF1 deadenylase, which prevents PyCCR4-1 association with the complex, leads to a similar phenotype. Comparative RNA-seq revealed that PyCCR4-1 affects transcripts important for transmission-related functions that are associated with male or female gametocytes, some of which directly associate with the immunoprecipitated complex. Finally, circular RT-PCR of one of the bound, dysregulated transcripts showed that PyCCR4-1 does not have gross changes in UTR or poly(A) tail length. We conclude that general and transmission-specialized members of the CAF1/CCR4/NOT complex play critical and intertwined roles in gametocyte maturation and transmission.

**AUTHOR SUMMARY:** Malaria is a disease caused by *Plasmodium* parasites, which are transmitted during an infectious blood meal by anopheline mosquitoes. Transmission of the sexual stages of the parasite to mosquitoes requires the proper regulation of specific mRNAs. While much work has been done to characterize regulation of mRNAs in female gametocytes, little has been done to assess this regulation in male gametocytes. Here, we demonstrate that PyCCR4-1, a member of the CAF1/CCR4/NOT RNA metabolic complex, acts upon transcripts both directly and indirectly in both male and female parasites, and results in a reduction of male gametocytemia. In gametocytes lacking PyCCR4-1, as well as those expressing a catalytically dead variant, the initial coordinated wave of male gametocyte activation is lost, and these parasites are less able to productively infect mosquitoes. We find that PyCCR4-1 requires its association with PyCAF1 and by proxy, the rest of the complex, in order to perform its functions based upon experiments in both *Plasmodium yoelii* and *Plasmodium falciparum*. We also find that the CAF1/CCR4/NOT complex is directly binding some of these transcripts and is likely acting both directly and indirectly to modulate transcript abundance. These findings demonstrate that the combined effects of the CAF1/CCR4/NOT complex upon specific mRNAs are important for both male and female gametocytes, and that this regulation is required for efficient transmission to the mosquito vector.

## INTRODUCTION

Malaria remains one of the great global health problems today, with 216 million new infections and 445,000 deaths attributed to it annually (1). Resistance to frontline drugs is spreading, and understanding the development and transmission of the malaria parasite is important to bolster efforts to reduce or eliminate deaths due to this infection. For the parasite to transmit from a vertebrate host to the mosquito vector, a small percentage of the cells will differentiate from asexual forms and develop into sexual stage gametocytes, which can persist in an infectious state until a mosquito takes a blood meal. This event allows a small number of gametocytes to be taken up into the mosquito, but with far fewer parasites productively infecting it (2). Following two weeks of development within the mosquito, a small number of sporozoites will similarly be injected into a host by the mosquito as it takes another blood meal (3). In the effort to develop vaccines and drugs, transmission events have been identified as prime targets because they are population bottlenecks in the parasite life cycle. We and others have focused upon the transmitted gametocyte and sporozoite stages of *Plasmodium* parasites to identify and exploit their weaknesses. In both cases, very few parasites are transmitted, and thus these bottlenecks are excellent points of intervention. The identification of molecular processes that are important for the transmission of the parasite in one or both of these events, and their modes-of-action, are thus top priorities for the development of new therapeutics.

Recent work has shown that the parasite requires tight transcriptional and translational control to navigate these complex transmission events (4–8). Despite the complex events required for the effective transmission of the parasite, *Plasmodium* has only one known, expanded family of specific transcription factors, the ApiAP2 proteins (reviewed in (9)). In *Plasmodium*, other ways to regulate gene expression that act post-transcriptionally to control translation, and to stabilize or degrade RNA, thus have increased importance (5, 10). For instance, the parasite utilizes RNA-binding proteins (e.g. DOZI, CITH) to impose translational repression and mRNA stabilization on transcripts in female gametocytes that are produced long before they are needed to establish a new infection of a mosquito (10–15). DOZI (<u>D</u>evelopment <u>o</u>f <u>Z</u>ygote <u>I</u>nhibited) is a DDX6 RNA helicase that is homologous with yeast DHH1 and human rck/p54, and CITH is a Lsm14 homolog of worm <u>C</u>AR-<u>I</u> and fly <u>T</u>railer <u>H</u>itch. A current model invokes these controls as a means for the parasite to always be ready to respond to external stimuli that indicate that transmission has occurred, and thus enables the rapid translation of the preserved mRNAs and establishment of the new infection (16). Moreover, this molecular process is essential to the parasite, as genetic deletion of *dozi* or *cith* results in a complete halt of development in early mosquito stage (13, 14). Similar regulatory events occur in the other transmitted stage (sporozoites) via the PUF2 RNA-binding protein, as deletion of *puf2* results in the gradual loss of infectivity and subsequent premature dedifferentiation into a liver stage-like form while in the salivary gland (7, 8, 17). As in model eukaryotes, many of these regulatory functions of RNA metabolism occur in cytosolic granules within the parasite as well (13, 14, 17, 18).

In addition to transcript stabilization, translational control can also be accomplished by degrading specific transcripts in a process typically initiated by deadenylases. This is accomplished by the degradation of the poly(A) tail that regulates the stability of the mRNA. Shortening the poly(A) tail to a critical length in turn promotes the subsequent decapping and complete degradation of the transcript by other factors (19, 20). In model eukaryotes, the main complex responsible for deadenylation is the CAF1/CCR4/NOT complex, which also participates in transcriptional elongation, translational repression, and histone modification functions, and thus acts broadly upon gene expression (20). This complex typically contains two putative deadenylases, with CAF1 (CCR4-Associated Factor 1) serving as the major deadenylase and CCR4 (Carbon Catabolite Repressor 1) playing additional or specialized roles, except for in yeast where the roles are reversed (20). While CAF1’s role in binding to and degrading poly(A) tracts is best appreciated, it has been shown to bind several other poly-nucleotide tracts (21). Recently, a cryo EM structure of the *S. pombe* CAF1/CCR4/NOT complex was reconstructed using immunoprecipitated material. This work confirmed previous studies that used recombinant proteins and binding assays to show that the complex is L-shaped, that NOT1 (Negative on TATA-less) acts as the scaffold, and that CCR4 binds to the complex indirectly through bridging interactions with CAF1 (21–23). While these associations and activities have been well described in model eukaryotes, little is known about the CAF1/CCR4/NOT complex’s form and function in malaria parasites.

In *Plasmodium*, previous work confirmed that normal deadenylase activity provided by CAF1 is essential for asexual blood stage growth (24, 25). Interestingly, insertion of a *piggyBac* transposon into the coding sequence revealed that CAF1 contributes to the regulation of invasion and egress-related genes in asexual blood stage parasites (24). It is possible that this transposon insertion still results in the production of a partially functional CAF1 protein. Multiple independent attempts to knock out *caf1* in the rodent-infectious species *P. berghei* failed, indirectly indicating that it is essential for parasite development (24, 26). Moreover, previous work on *Plasmodium* sporozoites identified that deletion of *pypuf2* led to significant changes in the transcript abundance of several members of the CAF1/CCR4/NOT complex (6, 27). Among the affected transcripts, two mRNAs encoding CCR4 domain-containing proteins were dysregulated. As the deadenylase proteins of the CAF1/CCR4/NOT complex have been shown to be specialized regulators in other species, we investigated the possibility that CCR4 domain-containing proteins may be acting in this capacity in *Plasmodium* as well (28, 29).

Here, we demonstrate that CCR4-1 is a specialized regulator during gametocytogenesis and transmission of the rodent-infectious *Plasmodium yoelii* parasite from the mammalian host to the mosquito vector. Deletion of *pyccr4-1*^-^, or expression of a putatively catalytic dead variant, resulted in a loss of the initial synchronous development of male gametocytes that can activate into gametes, as well as a reduction in the total number of mature male gametocytes.

Moreover, deletion of *pyccr4-1* also reduced the transmissibility of the parasite to the mosquito on both peak and post-peak transmission days, indicating that PyCCR4-1’s functions extend beyond its role in the synchronization of gametocytes. Comparative transcriptomics of wild-type and *ccr4-1*^-^gametocytes revealed that PyCCR4-1 significantly impacts the abundance of transcripts that are translationally repressed in female gametocytes, and those that impact the transmission to and establishment of an infection in the mosquito. We found that PyCCR4-1 binds directly to some of these affected transcripts and allows for increased transcript abundance without affecting UTR or poly(A) tail length. Surprisingly, this effect runs counter to the major canonical role of a deadenylase. Finally, proteomic characterizations and genetic modifications of *pycaf1* and *pfcaf1* indicate that PyCCR4-1 must associate with the canonical CAF1/CCR4/NOT complex in order to properly regulate transcripts associated with gametocyte development and to promote host-to-vector transmission.

## RESULTS

### PyCCR4-1 localizes to discrete cytosolic granules

In model eukaryotes, CCR4 performs many of its functions while in association with the other members of the CAF1/CCR4/NOT complex and is found in nuclear and cytosolic granular structures (19). Using immunofluorescence and live fluorescence assays with transgenic PyCCR4-1::GFP parasites, we observed that PyCCR4-1 localized to cytoplasmic puncta in asexual blood stage parasites, and is similarly localized in both male and female gametocytes (Figure 1, Fig S1AB). Moreover, this expression profile extends to oocysts, oocyst sporozoites, and salivary gland sporozoites, where PyCCR4-1 was seen both in cytosolic puncta and located diffusely throughout the parasite (Fig S1CD). However, PyCCR4-1 was not detected above background in liver stage parasites. Thus, the near constitutive expression and localization of PyCCR4-1 in cytoplasmic foci in *Plasmodium* resembles that of its orthologues in model eukaryotes.

**Figure 1:**
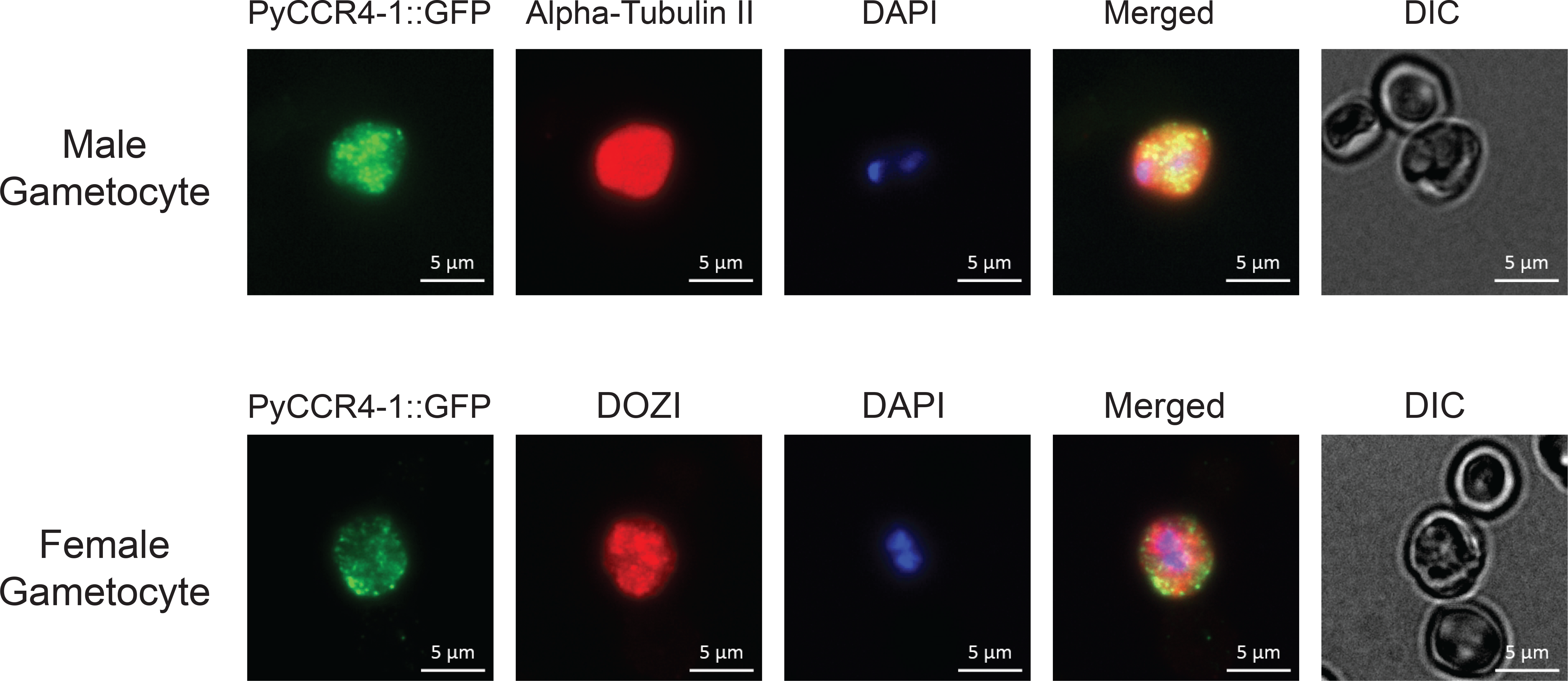
PyCCR4-1::GFP is expressed in cytosolic granules in sexual stage parasites. Representative images of sexual stages treated with DAPI and antibodies to GFP (to detect PyCCR4-1::GFP) or to stage-specific cellular markers (alpha-tubulin, or human DDX6 that cross-reacts with DOZI) are shown. Scale bars are 5 microns.

### PyCCR4-1 associates with a canonical CAF1/CCR4/NOT complex

Bioinformatically, we identified the genes for all members of the canonical CAF1/CCR4/NOT complex in *Plasmodium*, except for *not3* and *caf130*. The absence of these two particular genes is not surprising, as these genes are also absent in humans and *Drosophila* (19). In addition, we(Figure S2A). Total proteomics of mixed blood stage samples of *P. yoelii* wild-type and *pyccr4-1*^-^ parasite lines indicated that many of these bioinformatically defined members (NOT1, NOT4, NOT5, NOT Family Protein, CAF40) are expressed at levels that permit their detection, whereas CCR4-1 and CAF1 were not sufficiently abundant to be detected using highly stringent thresholds (Table S1).

To experimentally determine the composition of this complex in *Plasmodium yoelii*, the C-terminal GFP tag on PyCCR4-1::GFP was utilized to immunoprecipitate the CAF1/CCR4/NOT complex from synchronized schizonts, when PyCCR4-1 is most abundant and is most prominently localized to cytoplasmic granules. As seen in model organisms, PyCCR4-1 associates, directly or indirectly, with most members of the canonical CAF1/CCR4/NOT complex in *P. yoelii* (Table 1, Table S2) (20). Specifically, we found that PyCCR4-1 associates with CAF1, NOT1, CAF40, NOT2 and a NOT family protein above our most stringent SAINT (Significance Analysis of INTeractome) threshold (0.1), and with NOT5 using a less stringent threshold (0.1 to 0.35). A small number of peptide spectral matches for NOT4 were also observed, but were not sufficiently enriched to be confidently included. This low abundance of NOT4 is consistent with its known transient association with the CAF1/CCR4/NOT complex in other eukaryotes (30). We also found that PyCCR4-1 interacts with proteins involved in the nuclear pore complex and RNA export (e.g. karyopherin-beta 3, exportin-1, UAP56), proteins involved in translation initiation (e.g. eIF2A, EF-1, EIF3D, PABP), and translational repression (e.g. CELF2/Bruno, DOZI, CITH, PABP) (Table 1, Table S2) (20). All of these interactions are consistent with appreciated CAF1/CCR4/NOT interactions in model eukaryotes. Recently, a proteome of stress granule components in *S. cerevisiae* defined several cytosolic granule regulators, several of which we also found associated with PyCCR4-1 (31). Specifically, we identified that multiple CCT proteins of the TRiC complex (e.g. CCT4, CCT5, CCT8) and HSP40-A associate (SAINT score < 0.1), and this list expands to include the remainder of the TRiC core complex(e.g. TCP1, CCT2, CCT3, CCT6, CCT7) and a regulatory kinase (CK1) (SAINT scores between 0.1 to 0.35). In addition, recent work has implicated karyopherins/nuclear import receptors in the regulation of proteins found within liquid-liquid phase separations/cytosolic granules (32). Here, we identified that karyopherin beta 3 associates with the *P. yoelii* CAF1/CCR4/NOT complex, and perhaps indicates that similar regulatory processes are at work. These data indicate that the composition of the CAF1/CCR4/NOT complex, including the presence of cytosolic granule regulators, are likely conserved from model eukaryotes to *Plasmodium*.

**Table 1:**
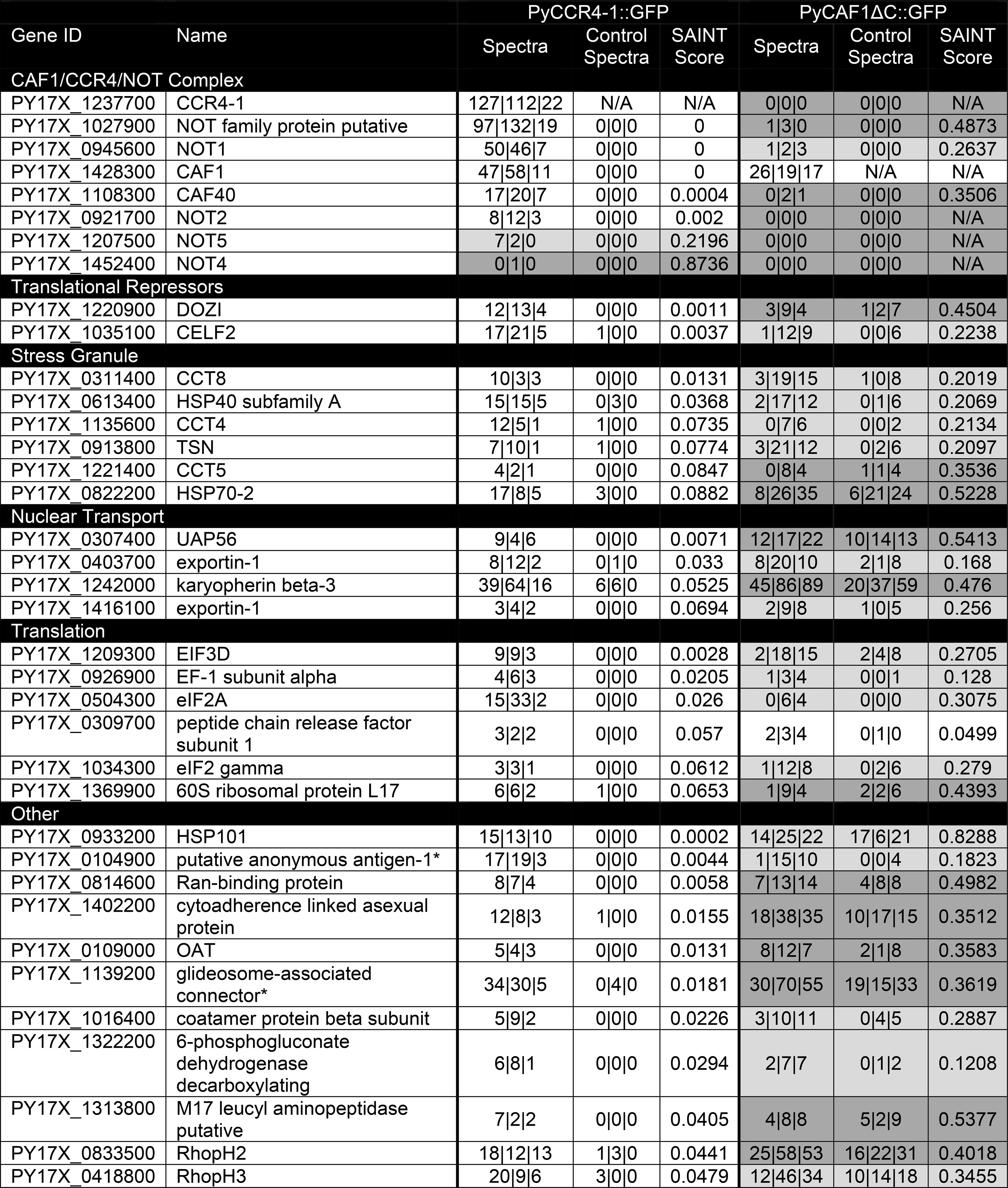
Interacting proteins with PyCCR4-1::GFP 200 and PyCAF1D::GFP proteins. Identified proteins are categorized based upon known or putative functions. Those that are currently unannotated are marked with an asterisk and described with the name of the closest protein identified by BLASTp alignment and/or a Phyre2 search (70, 71). Within each category, proteins are listed from lowest to highest SAINT score based on the PyCCR4-1::GFP data. Most stringent SAINT Score is unshaded, SAINT scores that fall between 0.1 and 0.35 are shaded in light gray and above 0.35 are shaded in dark gray. Strength of interactions in the other categories differ highly between the two immunoprecipitations. Average P is the average P value of all three biological replicates and SAINT Score is SAINT’s representation of FDR. Total number of experimental and control spectra for each replicate are also shown for comparison.

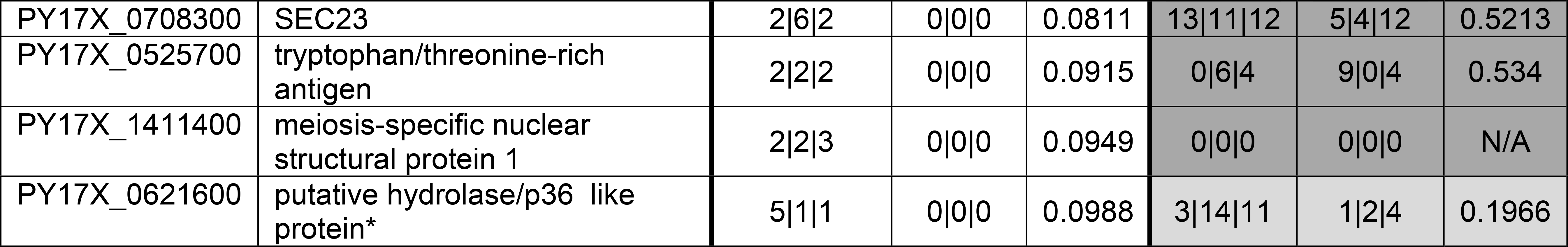

### PyCCR4-1 is important for the development and transmission of male gametocytes

As deadenylases are also known to act as translational regulators in specific and temporal manners, we investigated the role of all members of the CCR4 domain-containing protein family throughout the *Plasmodium* life cycle. Utilizing BLASTp alignments, we identified four high-confidence CCR4 domain-containing proteins in *Plasmodium* spp. that all have homology to deadenylases in other model eukaryotes (e.g. Yeast, Human, Mouse) (Figure S2AB). The domain architecture of CCR4-like proteins involves a Leucine Rich Repeat Region (LRR) and an Endonuclease/Exonuclease/Phosphatase (EEP) domain. The LRR mediates the interaction of CCR4 with CAF1 and the rest of the NOT complex, while the EEP domain contains active site residues required for deadenylation activity. Of these, we found that the EEP domain of PyCCR4-1 aligns most closely with the consensus CCR4 domain-containing proteins from model eukaryotes. However, beyond the CCR4-EEP domain, there is no significant homology between other regions from PyCCR4-1, 2, 3, and 4 to each other, or to homologues from model species (Figure S2B) (33). While a canonical LRR was not bioinformatically detectable in any of the CCR4 domain-containing proteins, immunoprecipitation of PyCCR4-1::GFP demonstrated that it retains the ability to associate with its complex (Table 1).

As our recent RNA-sequencing data from *Plasmodium yoelii* shows that all four genes are expressed in asexual blood stages and in gametocytes (4), we sought to determine if any of the CCR4 domain-containing proteins played an important, stage-specific role in the parasite life cycle. To this end, we replaced their coding sequences with a GFP-expression cassette and a human dihydrofolate reductase (HsDHFR)-expression cassette via double homologous recombination in the *Plasmodium yoelii* 17XNL strain (Figures S2CDEF). These lines were cloned via limiting dilution prior to characterization and their transgenic genotypes were confirmed using PCR across both homology regions. These clonal parasites revealed that deletion of any one of these genes individually was not lethal in asexual blood stages.

As CCR4 proteins are known to play specialized roles in eukaryotes, transgenic parasites lacking one of the four genes (*pyccr4-1, pyccr4-2, pyccr4-3, pyccr4-4*) were also compared to wild-type parasites in all stages of the parasite’s life cycle. Phenotypic observations were performed by measuring parasite numbers, prevalence of mosquito infection, and developmental timing/completion throughout the *Plasmodium* life cycle (Figure 2, Figure S3, Table S3). Deletion of *pyccr4-2, pyccr4-3*, or *pyccr4-4* resulted in transgenic parasites that behaved as wild-type in all life cycle stages (Table S3).

**Figure 2:**
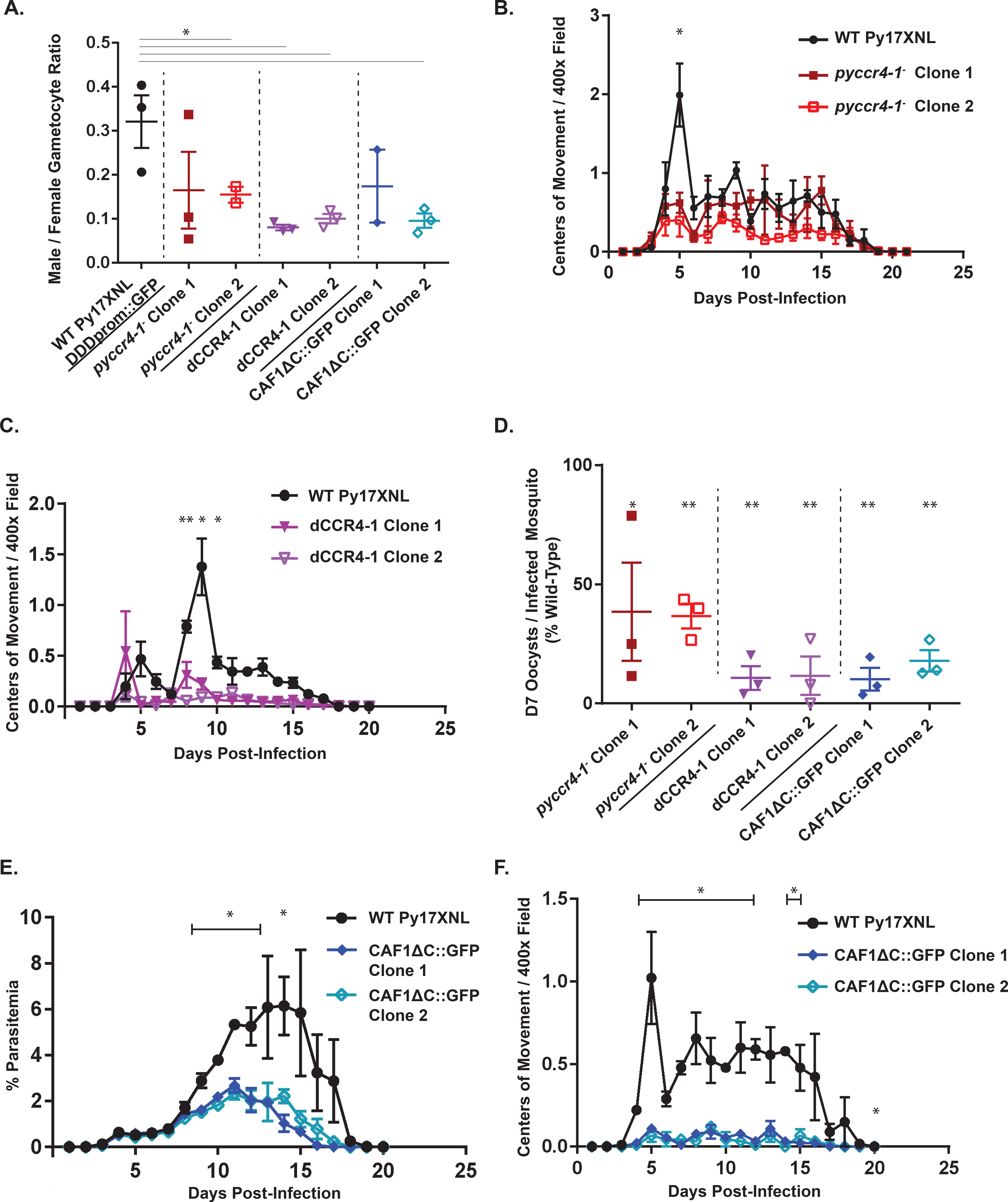
In *pyccr4-1*^-^, dCCR4-1, and PyCAF1ΔC parasites, coordination of gametocyte activation is lost and male gametocyte development and parasite transmissibility is reduced. A) The number of mature male gametocytes was determined by flow cytometry of sulfadiazine-treated and DDD/BIP-stained *P. yoelii* parasites. B,C,F) The number of centers-of-movement/exflagellation centers were quantified daily by microscopy on a 400× field upon injection with 10,000 (B, F) or 1,000 (C) blood stage parasites. Fewer WT and dCCR4-1 parasites were injected to better preserve animal health over the course of the experiment in accordance with our IACUC protocol (C). Plotted are three biological replicates with three technical replicates each. Error bars represent the standard error of the mean. D) The number of oocysts per infected mosquito on day seven post-infectious blood meal are plotted. Data represents at least 20 dissected mosquitoes per biological replicate conducted in triplicate. Error bars represent the standard error of the mean. E) Parasitemia was measured microscopically by giemsa-stained thin blood smears. Plotted are three biological replicates with three technical replicates each. Error bars represent the standard error of the mean.

However, while deletion of *pyccr4-1* had no effect upon asexual blood stage growth (Figure S3A), it led to significant effects during male gametocyte maturation and host-to-vector transmission (Figure 2). First, to assess gametocytogenesis and the number of mature male gametocytes present, we developed an antibody-based flow cytometry assay based in part upon the effective reporter system (820cl1m1cl1) commonly used in *P. berghei* (14). We generated antibodies against a recombinant domain variant of dynein heavy chain delta (PyDD, PY17X_0418900, “PyDDD” = AA1845-2334), and together anti-PvBiP antibodies to counterstain cells containing a parasite, we confirmed by flow cytometry and giemsa staining that PyDD is a marker for mature male gametocytes in *P. yoelii*, as it is in *P. berghei* This was further validated using a transgenic parasite line with a PyDDprom::GFPmut2 cassette integrated in the *p230p* safe harbor locus, where the population positive for both anti-PyDD and anti-GFP signals highly overlapped (Figure S3B). Ultimately, this approach allows these measurements to be done without the need to conduct reverse genetics in a base fluorescent reporter line, and frees GFP and RFP for other purposes.

Using this flow cytometric method, we found that transgenic *pyccr4-1*^-^ parasites produce fewer mature male gametocytes compared to wild-type parasites (Figure 2B, Figure S3D). Secondly, in contrast to wild-type parasites that have a semi-synchronous wave of gametocyte development, *pyccr4-1*^-^ parasites lose this coordination and instead develop fewer male gametocytes that can form gametes, and do so in an asynchronous manner (Figure 2B).

### The putative catalytic residues of PyCCR4-1 are required for its roles in gametocytogenesis and transmission

CCR4 proteins have well defined, conserved catalytic residues in model eukaryotes that are also conserved in *Plasmodium* species (Figure S2B) (34). To determine if the putative active site residues contribute to PyCCR4-1’s functions in male gametocytes, we created transgenic parasites with alanine substituted for two of the putative catalytic residues (D1852A, H1898A) of PyCCR4-1 (dCCR4-1) (Figure S3C). Like *pyccr4-1*^-^ parasites, dCCR4-1 transgenic parasites also produce fewer mature male gametocytes, and also lacked a synchronous wave of male activation (Figure 2 AC).

Because some male gametocytes retained the ability to mature and become exflagellating gametes in both the *pyccr4-1*^-^ and dCCR4-1 lines, we assessed whether they were transmissible to mosquitoes. In both transgenic lines, we observed a corresponding decrease of similar scale in the number of day seven oocysts compared to wild-type parasites when transmitted to *An*. *stephensi* on the peak day of male gametocyte activation into gametes (e.g. day five postinfection) (Figure 2D). Moreover, although there is no statistical difference in the number of male gametocytes that can activate between wild-type and *pyccr4-1*^-^ parasites after the peak day (Figure 2B, days six and beyond), a significant decrease (p<0.05) in the number of oocysts in the mosquito was still observed when parasites were transmitted two days post-peak (Figure S3E). These data indicate that the catalytic residues of PyCCR4-1 are required for normal male gametocyte development and host-to-vector transmission.

### Truncation of PyCAF1 prevents full assembly of the CAF1/CCR4/NOT complex and phenocopies the deletion of *pyccr4-1*

We next sought to determine if the typical association of PyCCR4 with the rest of its complex was required for its role in gametocyte development and host-to-vector transmission (35–37). As CCR4 domain-containing proteins associate with the NOT1 scaffold of the CAF1/CCR4/NOT complex indirectly by binding CAF1, genetic deletion of *caf1* would theoretically dissociate CCR4-1 from its complex. However, complete deletions of the *caf1* gene have been unsuccessful in both a conventional targeted attempt and in the PlasmoGEM broad-scale genetic screen in *P. berghei*, indicating that it is likely essential (24, 38). We have also attempted to completely delete the *P. yoelii caf1* coding sequence and similarly were unable to delete these sequences (Figure S4A). Instead, as the insertion of the *piggyBac* transposon into the *P. falciparum caf1* gene occurred in the coding sequence downstream of the CAF1 domain, we hypothesized that this portion of PfCAF1 may still be expressed and may be necessary (24). In support of this hypothesis, transcript expression analysis of the *P. falciparum* CAF1 disruptant line (PfCAF1ΔC) bearing this transposon insertion indicated that the CAF1 domain was still transcribed up to the insertion site, but not after (Figure S4B).

Based upon these expression data, we created a *Plasmodium yoelii* transgenic line that mimics this transposon insertion by inserting a C-terminal GFP tag and stop codon in the *Plasmodium yoelii caf1* gene in a comparable location following the CAF1 domain, thus creating a PyCAF1ΔC (AA 1-335) variant (Figure S4C) (24). We found that expression of the PyCAF1ΔC::GFP variant resulted in viable parasites, but importantly, that these parasites exhibit a similar growth attenuation as was observed for the *P. falciparum* PfCAF1ΔC line (Figure 2E) (24). To further assess the impact of the PyCAF1ΔC variant upon parasite growth and transmission, we observed comparable, but more pronounced, effects upon the activation of male exflagellation and parasite transmission as was seen with *pyccr4-1*^-^ parasites (10-fold decrease in male activation on peak day and >4-fold reduction in transmission to mosquitoes, respectively) (Figure 2F). These exacerbated effects may be caused by the combined effects of a reduction in total parasite numbers due to the deletion of portions of PyCAF1 and a PyCCR4-1-dependent defect in male gametocyte development.

To determine if the PfCAF1ΔC variant in human-infectious *P. falciparum* similarly impairs gametocytogenesis as was seen in rodent-infectious *P. yoelii*, the PfCAF1ΔC *piggyBac-insertion* parasite line was assessed for effects upon parasitemia, gametocytogenesis, as well as male gametocyte activation. The PfCAF1ΔC line exhibited significant decreases in gametocyte conversion, total gametocytemia, and exflagellation on the peak day as compared to wild-type *P. falciparum* NF54 strain parasites (Figure 3A-D, Table S4). These data support the observed *P. yoelii* phenotype and indicate that this conserved complex is important to sexual development across *Plasmodium* species.

**Figure 3:**
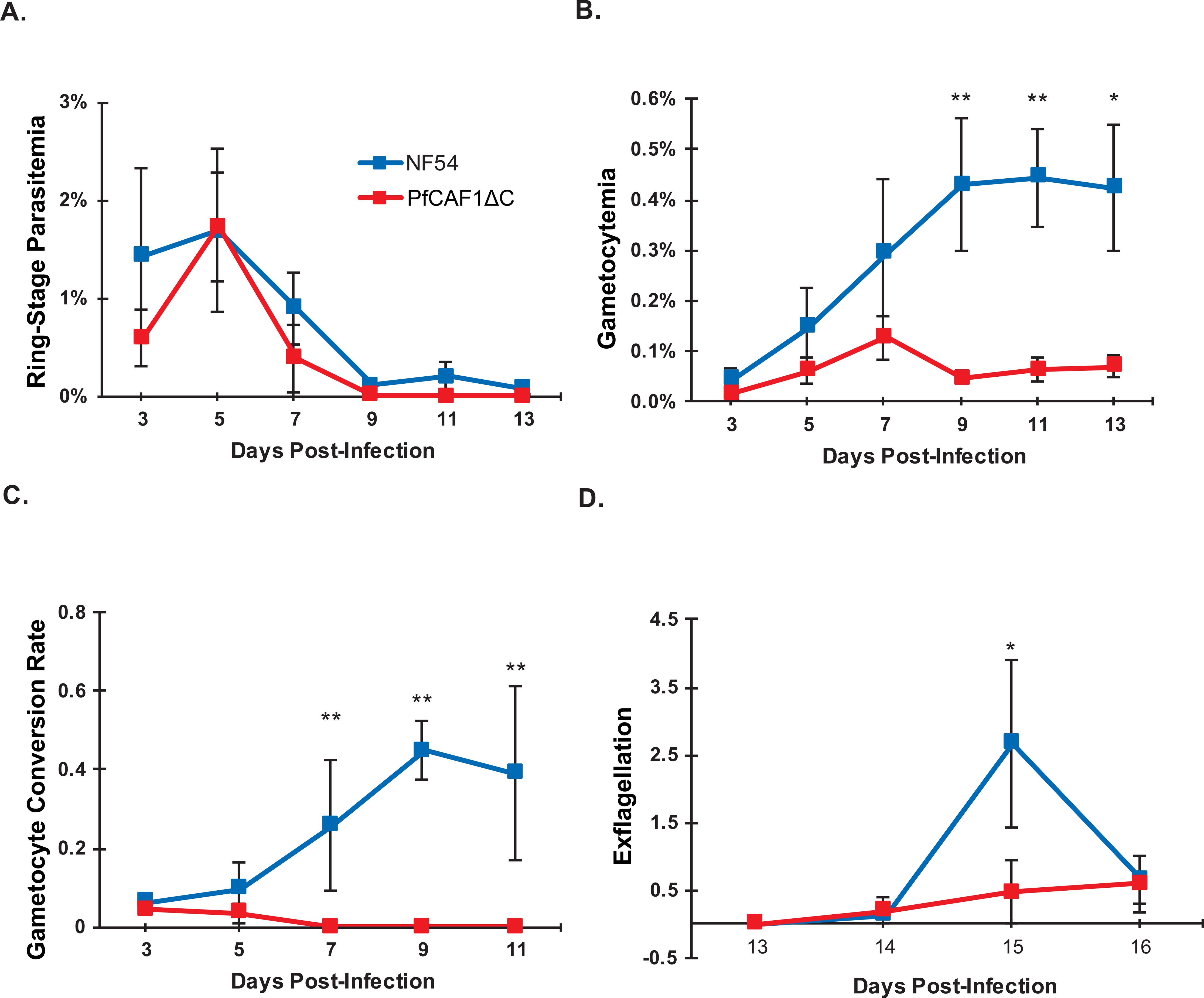
PfCAF1ΔC has decreased gametocyte conversion and exflagellation compared to wild type NF54 parasites. A) *P. falciparum* ring stage parasitemia and B) total gametocytemia counted in 10,000 RBCs averaged over a minimum of two biological replicates. Ring stages were used to represent asexual parasitemia as they were most easily distinguishable from dead/dying asexual forms. C) Conversion rates were calculated as described previously (56) by taking the stage II-gametocytemia on Day T and dividing by ring stage parasitemia on Day T-2. D) Exflagellation events were counted under 40× magnification for 10 fields of view. Error bars represent standard error of the mean. Statistical differences between wild-type and PfCAF1ΔC parasites were assessed via a paired Wilcoxon test. * = p<0.05, ** = p<0.01)

As we hypothesized that expression of only the CAF1 domain would prevent proper assembly of the CAF1/CCR4/NOT complex, we used IP-MS to determine if PyCAF1ΔC::GFP can still associate normally with its binding partners. We found that when *pycaf1* is disrupted in this manner, the resulting PyCAF1ΔC protein does not associate with most of the components of the CAF1/CCR4/NOT complex, including PyCCR4-1 (Table 1, Table S5). Utilizing the same stringent SAINT score (<0.1) for each immunoprecipitation, we found that only two proteins (PyCAF1ΔC itself, and subunit one of peptide chain release factor) are detected (Figure S5, Table S5). By expanding the SAINT threshold to 0.35, only 57 (41%) of PyCCR4-1::GFP’s 139 protein interactions were detected. Importantly, we did not detect any association of PyCAF1ΔC::GFP with PyCCR4-1 (no peptide spectral matches) and observed only a greatly reduced association with NOT1. These data suggest that PyCAF1ΔC is no longer able to interact with PyCCR4-1, while only weakly associating with other members of the CAF1/CCR4/NOT complex. This dysregulation is further observed in both asexual and sexual stage parasites, as full-length PyCAF1::GFP was found localized in cytosolic puncta, whereas PyCAF1ΔC::GFP was diffusely localized (Figure S6).

These changes in complex assembly caused by expression of the truncated PyCAF1ΔC variant, which phenocopies the deletion of *pyccr4-1*, led us to several conclusions. First, only the CAF1 domain of PfCAF1 and PyCAF1 is essential for parasite viability. Second, while portions of PfCAF1 and PyCAF1 C-terminal to the CAF1 domain are dispensable, they are critical for CAF1’s association with its complex, and essential for recruiting CCR4-1. Finally, these data indirectly suggest that PyCCR4-1 requires its association with PyCAF1 and the NOT complex in order to promote coordinated gametocytogenesis and efficient transmission to the mosquito.

### PyCCR4-1 affects important gametocyte and mosquito stage transcripts

Because PyCCR4-1 is a putative deadenylase, we hypothesized that these phenotypes in male and possibly female gametocytes may be attributed to PyCCR4-1 affecting the abundance of specific transcripts important to gametocytogenesis, gamete activation, and/or parasite transmission to mosquitoes. To determine the role of PyCCR4-1 in the regulation of transcripts in gametocytes, total comparative RNA sequencing (RNA-seq) was performed. Gametocytes from a wild-type line expressing GFP from the *p230p* dispensable locus (WT-GFP) and the *pyccr4-1*^-^ transgenic line were selected using sulfadiazine treatment, purified on an Accudenz gradient, and their RNA extracted for RNA-seq. Differential abundance of transcripts was assessed via DEseq2, and we utilized the p-adjusted value for all analyses (Figure 4A, Table S6) (39, 40).

**Figure 4:**
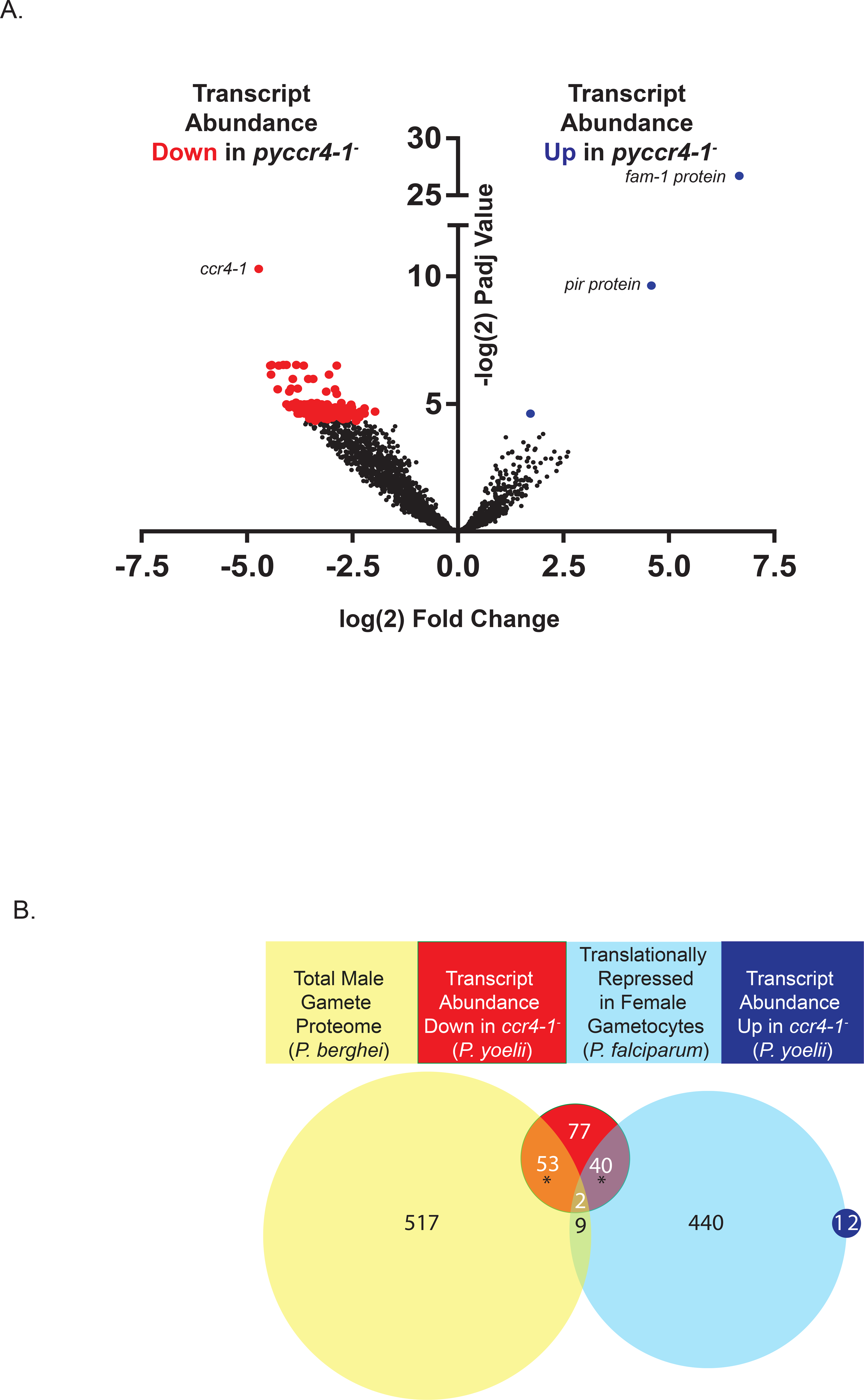
Transcripts with sex/transmission-related functions are modulated by PyCCR4-1. A) A volcano plot showing changes in transcript abundance up (blue), down (red) or unchanged (black). While few transcripts go up in abundance (from antigenically variant genes), nearly all affected transcripts decrease in abundance, thus indicating that PyCCR4-1 may play a role in preserving these mRNAs. B) In *pyccr4-1*^-^ gametocytes, twenty four percent of differentially abundant transcripts are translationally repressed in female gametocytes and another one third of the transcripts are enriched in the male gamete proteome (44, 66). * = p<0.01 by Fisher test

Nearly all (172 of 175) of the significantly affected transcripts (P-adjusted < 0.05, > 2 fold change) between WT-GFP and *pyccr4-1*^-^ parasites decreased in abundance in the *pyccr4-1*^-^ parasites, while only 3 transcripts increased in overall abundance (Table S6). Many of these decreases in transcript abundance are for 55 mRNAs that encode male-enriched proteins, and thus these changes can likely be attributed to the production of fewer mature male gametocytes in the *pyccr4-1*^-^ transgenic line. However, the effect upon other transcripts, including those associated with female gametocytes, cannot (Figure 4B). Most notably, transcripts that encode proteins involved in gamete function (e.g. GEST) and early mosquito stage development (e.g. p28, CITH, AP2-O, HMGB2, LAP2) decreased in abundance significantly in the absence of PyCCR4-1. Moreover, many of these transcripts are known to be translationally repressed in *P. falciparum* female gametocytes (Figure 4B). Interestingly, an ApiAP2 protein (PY17X_1417400) that is important for gene expression in gametocytogenesis decreased in abundance 10-fold in *pyccr4-1*^-^ gametocytes (41). Disruption of this ApiAP2 gene in *P. falciparum* by *piggyBac* transposon insertion resulted in the formation of no gametocytes (41). Other transcripts-of-interest that decreased in abundance are those that encode for multiple uncharacterized RNA-binding proteins (PY17X_1203900, PY17X_1457300, PY17X_0923600), a second ApiAP2 protein (PY17X_1317000) and BDP2 (PY17X_1431000), an uncharacterized putative transcriptional activator (Table S6). Together, we conclude that these differences in transcript abundance reflect both a reduction in the number of mature male gametocytes, but also indicate that PyCCR4-1 is acting to preserve specific transcripts important for the gametocyte and early mosquito stage parasite.

### The CAF1/CCR4/NOT complex specifically binds transcripts that are dysregulated in *pyccr4-1*^-^ parasites

As transcript abundances could be affected directly by PyCCR4-1 and its complex, or indirectly through compensatory mechanisms such as gene buffering when the *pyccr4-1* gene is deleted (42), we determined if these dysregulated mRNAs were bound by the CAF1/CCR4/NOT complex. To this end, unfused GFP (expressed in WT-GFP parasites) and PyCCR4-1::GFP were immunoprecipitated from purified, transgenic gametocytes, and the association of coprecipitated transcripts was detected by RT-PCR. We found that PyCCR4-1::GFP interacted specifically with a number of selected transcripts that substantially change in abundance in *pyccr4-1*^-^ parasites, including *p28* (PY17X_0515900), *lap2* (PY17X_1304300), and *nek3* (PY17X_0603200) (Figure 5A, top row). These transcripts are notable, as they are all important/essential for gametocytogenesis or transmission, and include transcripts known to be important to male (*nek3*) and/or female (*p28, lap2*) gametocytes (43–47). However, not all dysregulated transcripts (*cith* and *ap2-o*), nor unaffected housekeeping transcripts (*gapdh, eif2b*) were found specifically associated with PyCCR4-1 (Figure 5A, bottom row), suggesting that these effects likely result from a combination of both direct and indirect effects upon specific transcripts.

**Figure 5:**
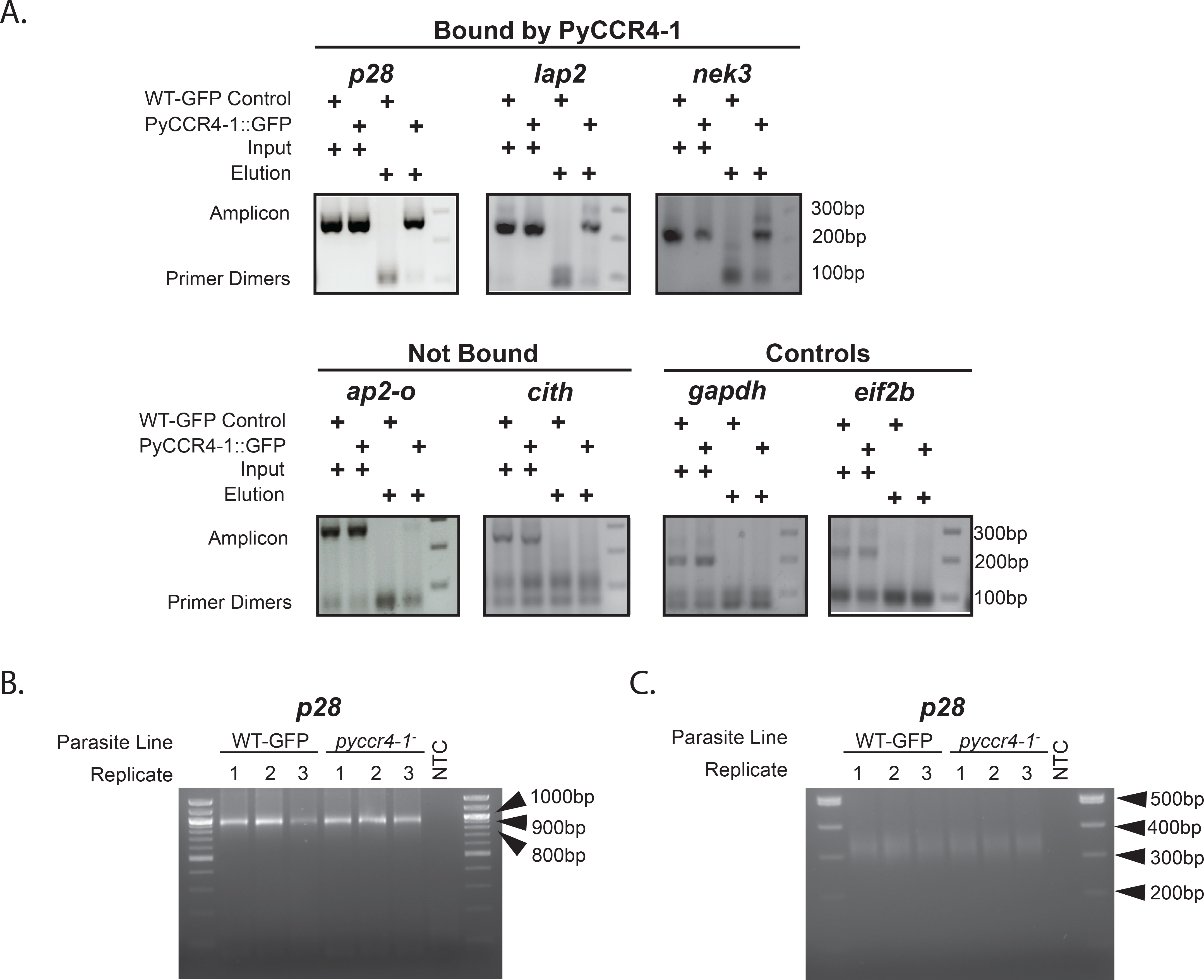
The CAF1/CCR4/NOT complex associates with some dysregulated transcripts but doesn’t grossly affect UTR/poly(A) tail length. A) Immunoprecipitation of PyCCR4-1::GFP allowed detection of the association of three selected transcripts that are affected by *pyccr4-1*^-^ (top), whereas other affected transcripts do not associate with PyCCR4-1::GFP (bottom, left). Two control transcripts that do not change upon deletion of *pyccr4-1* also do not interact with PyCCR4-1::GFP (bottom right). Shown are input and elution samples of a constitutive GFP-expressing clone and a PyCCR4-1::GFP clone. Amplicons and primer dimer bands are indicated with arrows. B and C) Circularized RT-PCR (cRT-PCR) of *p28* was conducted in both wild-type and *pyccr4-1*^-^ parasites to detect effects by PyCCR4-1 upon UTR and poly(A) tail length in purified gametocytes. Primers were designed within the coding sequences (B) or near the poly(A) tail (C) as defined by sequencing of cRT-PCR products from panel B, which allow observations of UTR and poly(A) tail lengths respectively. Three biological replicates and a No Template Control (NTC) are shown with NEB 100bp molecular weight ladder in parallel. Extended data is available in S7 and S8 Figures.

The direct effects that CCR4 can have on transcript abundance in model eukaryotes have resulted from deadenylation of a target transcript, or through translational repression by binding/tethering to its complex(28). To investigate this, circular-RT PCR (cRT PCR) was used to determine if the UTR and poly(A) length of both a control transcript (*gapdh*, not affected by *ccr4-1* deletion, does not interact with CCR4-1), and an affected/bound transcript (*p28*) were impacted by PyCCR4-1. cRT PCR demonstrated that, upon *pyccr4-1* deletion, there were no gross effects on UTR/poly(A) tail lengths of these transcripts using primers that anneal near the start and stop codons (Figure 5BC, oligonucleotides provided in Table S7). Sequencing of cloned PCR products from both wild-type and *pyccr4-1*^-^ samples revealed the consistent composition of the 5’ and 3’UTRs, as well as the presence of a poly(A) tail of the *p28* transcript (S8 Figure). However, none of these sequencing runs could resolve the precise length of the poly(A) tail, but did permit design of primers near to the poly(A) tail to assess the distribution of poly(A) tail lengths in the population. Using these primers, we did not observe any differences in poly(A) length between the wild-type and *pyccr4-1*^-^ populations (Figure 5C). These data indicate that the direct effect of PyCCR4-1 on these transcripts in gametocytes do not impact the poly(A) tail/UTR length suggesting that the complex is acting in other ways to preserve these transcripts.

## DISCUSSION

*Plasmodium* encodes few known specific transcription factors and a relatively over-represented number of RNA-binding proteins (10% of its predicted proteome) (27, 48). One model suggests that is has adapted these complementary regulatory mechanisms to achieve its preferred RNA homeostasis. Moreover, the malaria parasite also proactively transcribes a large number of genes before transmission, whose proteins are only required post-transmission. This translational repressive mechanism has been shown to be imposed by members of the DOZI/CITH/ALBA complex, as well as by PUF2. Here, we demonstrate that the PyCCR4-1 and PyCAF1 members of the CAF1/CCR4/NOT complex play additional roles in either the preservation or expression of translationally repressed transcripts through direct and indirect means (Figure 6). Moreover, we find that PyCCR4-1 is also important for the development of the male gametocyte, as well as for the efficient transmission of gametocytes to the mosquito vector.

**Figure 6:**
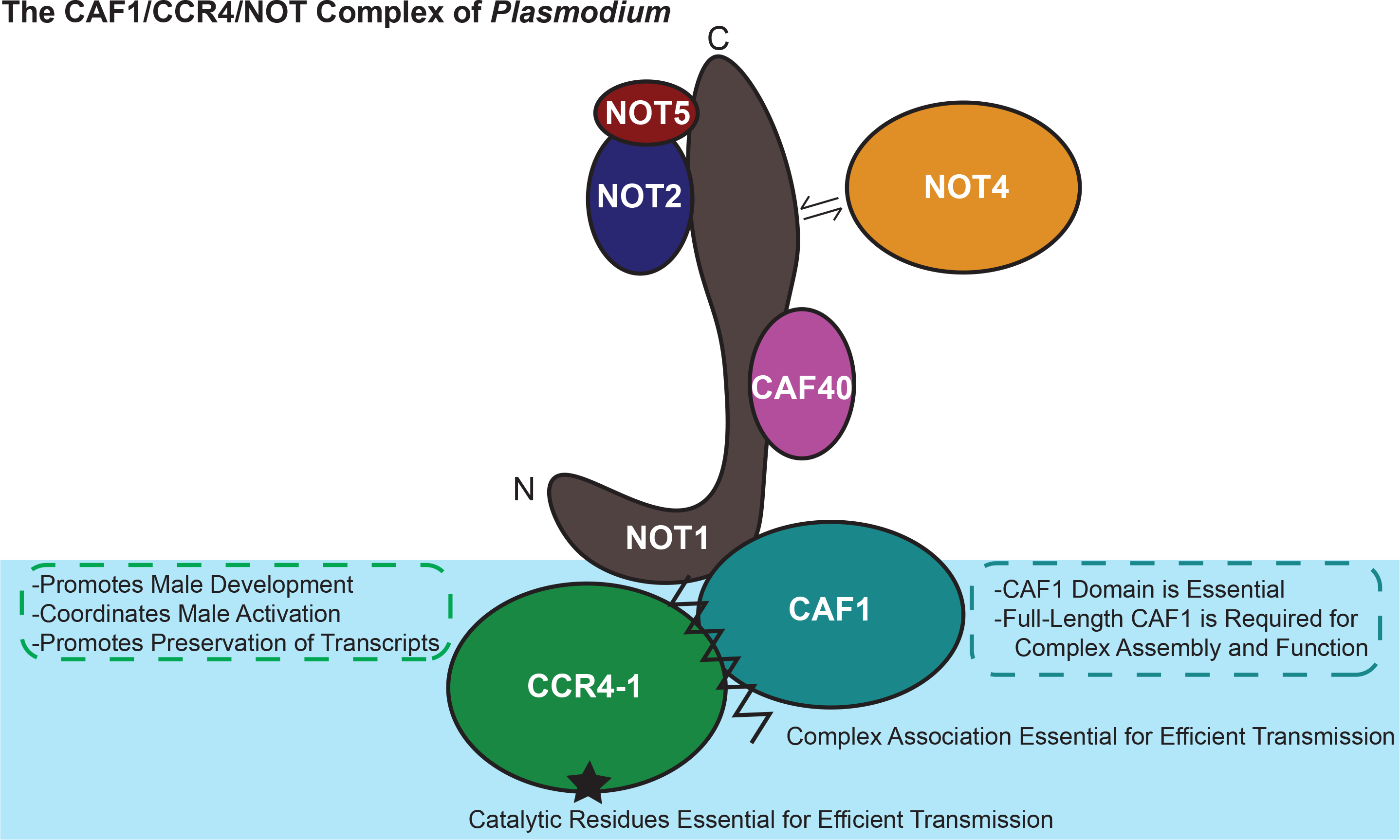
PyCCR4-1 acts as a regulator of specific transcripts in gametocytes while CAF1 acts more generally. The CAF1/CCR4/NOT complex in *Plasmodium* as identified through crosslinking IP-MS is shown. Contacts are inferred through previous studies in model organisms and sequence conservation of the interaction domains (19, 23, 67–69).

The composition and behaviors of the CAF1/CCR4/NOT complex are well-conserved in eukaryotes, but key species-specific differences exist. First, our proteomic and bioinformatic analyses of *Plasmodium* species showed that many of the core components of this complex are shared between *Drosophila melanogaster, Saccharomyces cerevisiae, and Homo sapiens* (NOT1, NOT2, NOT5 (NOT3), NOT4, CCR4, CAF1 (POP2), and CAF40), but that some proteins are apparently not encoded at all (CNOT10) (27). Second, while there are well-defined roles for this complex in both the nucleus and cytoplasm of model eukaryotes, we find by IFA that the vast majority of PyCCR4-1, PyCAF1, and by inference the entire complex, localizes to discrete cytosolic granules throughout most of the *Plasmodium* life cycle. For this reason, we have focused our analyses here upon the known cytoplasmic roles of the complex, but additional work to define possible nuclear functions of the CAF1/CCR4/NOT complex is certainly warranted. Third, in model eukaryotes CAF1 acts as a bridge to recruit CCR4 to the NOT1 scaffold. Interestingly, the CAF1 domain is typically required for the association of CAF1 with the NOT complex, which although it is the essential domain of the protein, is not sufficient for robust binding to its complex. Moreover, in *Plasmodium* the CAF1 domain alone does not permit recruitment of PyCCR4-1 to PyCAF1, indicating that sequences C-terminal of this domain in *Plasmodium* help mediate these interactions. In trypanosomes, the interaction of CAF1 with the rest of the NOT complex is affected by CNOT10, which is not bioinformatically identifiable in *Plasmodium* and thus may also explain why PyCAF1ΔC fails to properly assemble with its complex (42). This C-terminal truncation of CAF1 may result in the removal of the binding site of a yet-to-be-identified protein that fulfills this role. Finally, this exacerbated phenotype of the PyCAF1ΔC parasite line matches what has been previously seen in CAF1 mutants in yeast, perhaps due to its more central, coordinating position in the complex (49).

Previous studies of CCR4 in model eukaryotes have demonstrated that CCR4 can act as a translational repressor, and that its catalytic residues and deadenylase activity are implicated in its repressive functions (28, 50, 51). Our results demonstrate that PyCCR4-1 helps to preserve translationally repressed transcripts and is suggestive of similar roles in *Plasmodium* (Figure 4). Furthermore, transgenic parasites that lack *pyccr4-1*, that express a putatively dead catalytic variant of PyCCR4-1, or that express a truncated variant of PyCAF1 (PyCAF1ΔC) demonstrate the same phenotype, and thus we conclude that the catalytic residues of the PyCCR4-1 protein and its association with its complex are required for this regulatory function (Figure 2, 6). Additionally, as the maturation of male gametocytes is substantially promoted by the presence of catalytically active PyCCR4-1, it is unsurprising that one third of the transcripts that PyCCR4-1 affects encode proteins that are present in *a P. berghei* male gamete proteome (Figure 4). It should be noted that the PyCCR4-1 catalytic mutant (dCCR4-1) has a slightly stronger effect upon gametocytogenesis, gametogenesis, and parasite transmission than does the deletion of *pyccr4-1*. This effect could result if dCCR4-1 is still able to bind transcripts and prevent other proteins from assembling onto them, thus acting as a dominant negative variant.

Similarly, expression of a truncated CAF1 variant in *P. yoelii* or *P. falciparum* produced similar phenotypes during gametocytogenesis and male gametocyte activation (Figures 2 and 3). Proteomic and IFA analyses of PyCAF1ΔC showed that this could be explained in part because this variant no longer associated with PyCCR4-1, only weakly bound with the majority of the CAF1/CCR4/NOT complex, and became diffusely localized (Table 1, Figure S6). This work also defines the CAF1 domain as the essential portion of the PfCAF1 and PyCAF1 proteins, and that sequences C-terminal of it play roles in association with NOT1 and CCR4-1 and contribute to asexual parasite growth.

During sexual development we find that PyCCR4-1 binds to multiple transcripts essential for gametocyte development and host-to-vector transmission, which are dysregulated in *pyccr4-1*^-^, parasites. We hypothesize that PyCCR4-1 is interacting with and acting upon these transcripts in a concerted effort to preserve them for use post-transmission. Circular RT-PCR demonstrated that UTR and poly(A) lengths of an affected female transcript (*p28*) is not affected in *pyccr4-1*^-^ transgenic parasites. Therefore, we posit that PyCCR4-1 and its complex can be utilizing functions independent of deadenylation to achieve these ends. For instance, tethering of transcripts to the CAF1/CCR4/NOT complex is able to induce repression of a target transcript in other systems, even in the absence of deadenylation of those transcripts (28, 52). Taken together, these data indicate that the *Plasmodium* CAF1/CCR4/NOT complex provides key functions in the regulation of specific transcripts to promote coordinated male gametocyte maturation and parasite transmission.

## MATERIALS AND METHODS

Extended versions of materials and methods are provided in S1 File.

### Ethics Statement

All animal care strictly followed the Association for Assessment and Accreditation of Laboratory Animal Care (AAALAC) guidelines and was approved by the Pennsylvania State University Institutional Animal Care and Use Committee (IACUC# 42678-01). All procedures involving vertebrate animals were conducted in strict accordance with the recommendations in the Guide for Care and Use of Laboratory Animals of the National Institutes of Health with approved Office for Laboratory Animal Welfare (OLAW) assurance.

### Experimental Animals

Six-to-eight week old female Swiss Webster mice from Harlan (recently acquired by Envigo) were used for all of the experiments in this work. *Anopheles stephensi* mosquitoes (obtained from the Center for Infectious Disease Research; Seattle, WA) were reared at 24C and 70% humidity and were used to cycle *Plasmodium yoelii* (17XNL strain) parasites.

### Production of Transgenic Parasite Lines

Transgenic *Plasmodium yoelii* (17XNL strain) parasites were created using targeting sequences to incorporate sequence into the target gene using double homologous recombination using standard procedures (53). Parasite genomic DNA was purified (QIAamp DNA Blood Kit, Qiagen, Cat# 51106) and genotyping PCR was performed to assess the ratio of WT to transgenic parasites present. Clonal parasite populations were produced using limiting dilution cloning. The PfCAF1ΔC line was previously generated as described (24). Validation of CAF1 transcript expression was performed via RT-PCR on 100ng DNase-treated RNA from PfCAF1ΔC and NF54-control parasites using primer sets supplied in S7 Table.

### Production and Accudenz Purification of *P. yoelii* Schizonts and Gametocytes

To produce schizonts in culture, infected Swiss Webster mice were exsanguinated by cardiac puncture and the blood was collected into complete RPMI (cRPMI), spun at 200 *xg* for 8 min to remove the serum, and then cultured in 30 ml cRPMI in a 5% CO_2_, 10% O, 85% N gas mixture for 12 hours at 37C. Cultures were underlayed with 10 ml of 17% w/v Accudenz in 5 mM Tris-HCl (pH 7.5@RT), 3 mM KCl, 0.3mM disodium EDTA, 0.4× PBS (without calcium and magnesium) and spun at 200 *xg* for 20 min with no brake (53). Parasites were collected from the interface between the Accudenz and cRPMI layers, transferred to a fresh conical tube, supplemented with an equal volume of additional cRPMI, and spun for 10 min at 200 *xg*. The supernatant was then removed and the parasite pellet processed for downstream applications.

Gametocytes were produced by treatment of the mice at 1% parasitemia with 10 mg/L sulfadiazine (VWR, Cat# AAA12370-30) in their drinking water for two days prior to exsanguination. The blood was maintained in warm (37C) cRPMI to prevent activation of gametocytes and parasites were purified as described above.

### *P. falciparum* Gametocyte Production

Gametocyte-producing cultures were established as described previously (54) with some modification. Briefly, starter cultures of wild-type *P. falciparum* NF54 and PfCAF1ΔC were grown to ^~^5% parasitemia in standard culture conditions in cRPMI supplemented with 25 mM HEPES, 0.2% D-glucose, 200 uM hypoxanthine, 0.2% w/v sodium bicarbonate, and 10% v/v heat-inactivated human serum at 6% hematocrit in a tri-gas incubator (5% CO_2_, 5% O_2_) at 37C. On Day 0, starter cultures were then used to inoculate 75 cm^2^ flasks (15ml culture volume at 6% hematocrit) in technical duplicate for each line at 0.5% parasitemia. Parasites were cultured for 17 days with daily media changes and no fresh addition of blood. Samples were taken to monitor parasite development starting at Day 3 post-infection and then every 48 hours until Day 13 post-infection, determined through Giemsa-stained smears. At seven days postinfection, technical replicate flasks were combined into one flask, which was maintained for the duration of the experiment. Samples were assessed in biological triplicate.

### PyDD Recombinant Protein Expression and Purification

Coding sequence for a domain of PyDD (PY17X_0418900, “PyDDD” = AA1845-2334) was generated by IDT as a codon-optimized gene block (gBlock) for expression from a modified pET28b+ vector (pSL0220) in *E. coli* BL21 (DE3) pLysS CodonPlus bacteria. Protein was purified first by standard Ni-NTA and then glutathione resin approaches (details provided in S1 File). Purity was confirmed to be >90% by SDS-PAGE by Coomassie Blue staining. Antibodies were generated in rabbits (screened for pre-immune sera with minimal background reactivity) by Pocono Rabbit Farm and Laboratory (Canadensis, PA).

### Live Fluorescence and IFA Microscopy

PyCCR4-1 and PyCAF1 expression in blood stages, oocyst sporozoites, salivary gland sporozoites and liver stages was observed by an indirect immunofluorescence assay (IFA), and expression in day seven oocysts was observed by live fluorescence. All samples for IFA were prepared as previously described, with all details provided in S1 File (55). Fluorescence and DIC images were taken using a Zeiss fluorescence/phase contrast microscope (Zeiss Axioscope A1 with 8-bit AxioCam ICc1 camera) using a 40X or 100X oil objective and processed by Zen imaging software.

### Measurement of Blood Stage Growth Kinetics

Cryopreserved blood infected with either wild-type (Py17XNL), *pyccr4-1*^-^, dCCR4-1, or PyCAF1ΔC parasites were injected intraperitoneally into Swiss Webster starter mice and parasitemia was allowed to increase to 1%. This blood was extracted via cardiac puncture and diluted in RPMI to 10,000 parasites per 100 ul (CCR4-1, CAF1ΔC) or 1,000 parasites per 100ul microliter (dCCR4-1). One hundred microliters was injected intravenously (IV) into three mice per replicate for each parasite line. Three biological replicates were conducted, each with three technical replicates. Parasitemia was measured daily by giemsa-stained thin blood smears. Centers of movement/exflagellation centers were also measured daily via wet mount of the blood incubated at room temperature for 10 min by counting the number of exflagellating male gametocytes in a confluent monolayer per 400× field (40× objective × 10× eyepiece). *P. falciparum* ring stage parasitemia and total gametocytemia were calculated every two days starting on Day 3 post-infection by averaging counts in 10,000 RBCs across a minimum of two biological replicates (provided in S6 Table). Sexual conversion was calculated as described previously (56) by taking the stage II-gametocytemia on Day T and dividing by ring stage parasitemia on Day T-2. Samples were taken for exflagellation assays on days 13, 14, 15, and 16 post-infection. Two-hundred microliter samples were taken from each flask and spun down at 0.3 *xg* for 30 seconds. Supernatant was removed and a 20 ul aliquot of remaining blood pellet was mixed with 20 ul of heat-inactivated human serum previously warmed to 37°C. The mixture was then allowed to incubate at room temperature for 15 min, after which exflagellation events were counted under 40× magnification for 10 fields-of-view.

### Flow Cytometry Gametocyte Counts

Cryopreserved blood infected with either wild-type (Py17XNL), *ccr4-1*^-^, dCCR4-1, or CAF1AC parasites was injected intraperitoneally into starter mice and transferred as above (10,000 parasites/100ul). On Day 5, gametocytes were produced by treatment of the mice with 10 mg/L sulfadiazine (VWR, Cat# AAA12370-30) in their drinking water for two days. Blood was collected by cardiac puncture and maintained in warm cRPMI to prevent activation of gametocytes and spun at 37°C. Blood was then fixed, passed through a cellulose column and stained as described above for IFA. Parasites were stained with the following primary antibodies: mouse anti-PvBIP Clone 7C6B4 (1:1000; (57)) and rabbit anti-PyDynein Heavy Chain Delta Domain (“PyDDD”, PY17X_0418900 AA: 1845 to 2335)) (1:1000, Pocono Rabbit Farm & Laboratory, Custom PAb), along with goat anti-mouse conjugated to AF594 (Fisher Scientific, A11012) and goat anti-rabbit conjugated to AF647 (Fisher Scientific, PIA32733) secondary antibodies. These were then analyzed on a LSR Fortessa (BD) in tube mode and collected samples were analyzed in FlowJo.

### Mosquito Transmission Studies

Cryopreserved blood infected with either wild-type (Py17XNL), *ccr4-1*^-^, dCCR4-1, or CAF1AC parasites was injected intraperitoneally into starter mice and transferred as above. Centers of movement were checked daily as above and mice were fed to mosquitoes on the peak day of exflagellation (day 5). Mosquito midguts were dissected at D7 post feed and analyzed for the prevalence of infection and oocyst numbers by microscopy. Mosquito midguts (day 10) or salivary glands (day 14) were dissected, ground, and sporozoite numbers counted.

### Immunoprecipitations, Western Blotting, and Mass Spectrometric Proteomics

Parasite pellets (schizonts) were crosslinked in 1% v/v formaldehyde and lysed using RIPA lysis buffer with a 1× protease inhibitor cocktail and 0.5% v/v SUPERase In, dounce homogenization with a tight pestle, and sonication. The parasite lysate was then precleared using streptavidin-coated dynabeads was immunoprecipitated using a biotin-conjugated anti-GFP antibody loaded on streptavidin-coated dynabeads for three hours at 4C with rotation. The beads were washed with modified RIPA wash buffer (50 mM Tris-HCl (pH 8.0@RT), 1 mM EDTA, 150 mM NaCl, 1% v/v NP40) once and then transferred to a new tube. The beads were washed 3 more times with modified RIPA wash buffer and then eluted at 45C overnight in a heat block. Samples were quality controlled by western blotting, and then subjected to tryptic digest and LC/MS/MS identification (Harvard Proteomics Core, run parameters listed in S1 File). The data was processed using the Trans-Proteomic Pipeline (TPP) (58) as described previously with few modifications (17). Spectra were searched against reference sequences downloaded in February 2016 from *Plasmodium yoelii* 17X (PlasmoDB, v27), mouse (Uniprot), and common contaminants (Common repository of adventitious protein sequences, (59) and randomized decoys generated through TPP. iX!Tandem and Comet searches were combined in iProphet (60) and protein identifications were determined by Peptide Prophet. Only proteins with a highly stringent false positive error rate of less than 1% are reported. To combine replicate proteomics datasets, SAINT version 2.5.0 was used (61). Only proteins with SAINT scores below 0.1 (most stringent) or 0.35 (stringent) were considered significant hits and included in the analyses, as used previously (2, 4, 6). The total proteome of Py17XNL mixed blood stages was determined using the same workflow (Penn State Proteomics Core).

### Total and Comparative RNA-seq

Gametocytes were produced, collected, and purified by an Accudenz gradient, as above. Infected RBCs were lysed with saponin, washed with 1×PBS, and released parasites were then lysed immediately using the QIAgen RNeasy Kit using the manufacturer’s protocol with the additional on-column DNaseI digestion. RNA yields were quantified spectrophotometrically by NanoDrop, and RNA samples were further quality controlled (BioAnalyzer) and used to create barcoded libraries (Illumina TruSeq Stranded mRNA Library). An equimolar pool of all samples was made and 100 nt single end read sequencing was performed on an Illumina HiSeq 2500 in Rapid Run mode. The resulting data was mapped to the *P. yoelii* 17XNL strain reference genome (plasmodb.org, v32 using Tophat2 in a local Galaxy instance (version .9). Gene and transcript expression profiles for both WT-GFP and *ccr4-1*^-^ assemblies were generated using htseq-count (Galaxy version 0.6.1galaxy3) (62) using the union mode for read overlaps. Count files were merged and compared using DESeq2 (Galaxy version 2.11.39 (63)). Six biological replicates were used for the WT transcriptomic profile, while four replicates were used in for the *ccr4-1*^-^ profiles. These were analyzed by a mean fit type with outlier replacement turned on to normalize the variance between the count files. The P-adjusted value was used for all analyses.

### Circular Reverse Transcription PCR (cRT-PCR)

RNA was isolated from purified *P. yoelii* wild type or *pyccr4-1*^-^ gametocytes by TRIzol/chloroform extraction and extensive DNaseI digestion. The 7-methylguanosine cap was removed from 10ug of total RNA using 2.5U Cap-Clip Acid Pyrophoshatase in 1×Cap-Clip Buffer supplemented with 10U Murine RNase Inhibitor at 37C for 1 hour. Treated RNA was TRIzol extracted, precipitated, dried, and then circularized with T4 RNA Ligase in T4 DNA Ligase buffer supplemented with 10% w/v PEG8000 and 10U Murine RNase Inhibitor at 16C for 24 hours. RNA was purified, precipitated, dried, and then subjected to reverse transcription using SuperScript IV and gene-specific primers (S7 Table). Specific PCR amplification of *gapdh* and *p28* sequences from the resulting cDNA was conducted using Phusion polymerase (NEB) and gene specific primers (Supp Table 7).

### Statistical Analyse

Statistical differences between *P. yoelii* wild-type and transgenic parasites were assessed via a two-tailed t-test on Graphpad Prism. Statistical differences between *P. falciparum* wild-type and PfCAF1ΔC parasites were assessed via a paired Wilcoxon test using R v. 3.3.1 (64) with p < 0. 05 indicating statistical significance.

### Data Availability Statement

All data is publically available on common data repositories. Proteomics data is accessible at the ProteomeXchange Consortium via the PRIDE partner repository with the dataset identifier PXD007042 (65). Transcriptomics data (both RAW and processed files) is accessible at the GEO repository (Accession # GSE101484). Details of datasets and identifiers are available in S1 File.

## ACKNOWLEDGEMENTS

We thank the Penn State University and Harvard University mass spectrometry cores for proteomic analyses, the Penn State University genomics core for RNA-sequencing, and Penn State Animal Resources for care of our animals. We thank Joe Reese for the anti-DDX6 antibody and Istvan Albert for critical discussion of transcriptomics analysis methods. We thank the Lindner and Llinás labs for critical discussion of the manuscript and its data. We thank Manuel Llinas for discussion and critical review of the manuscript. We thank Alison Roth and Shulin Xu for discussion of *P. falciparum* gametocyte culturing methods. We thank Hannah Haines for help counting slides.

## SUPPLEMENTAL FIGURE AND TABLE LEGENDS

<u>S1 Figure (.PDF)</u>: Genotyping PCR of (A) *pyccr4-1::gfp* Insertion of a C-terminal GFP tag was created using double homologous recombination at the 3’ end of the *pyccr4-1* coding sequence. Genotyping was performed by PCR on parasites using the primers indicated (listed in S7 Table). Independent clones were analyzed with a Py17XNL wild-type control, a no template control, and a plasmid positive control in parallel. (B, C, D) PyCCR4-1::GFP is expressed in asexual and mosquito stage parasites but is not detectable in liver stage parasites. Representative images of (B) asexual blood stage parasites, (C) oocyst sporozoites, salivary gland sporozoites and (D) 24 hour and 48 hour liver stage parasites treated with DAPI and antibodies to GFP (to detect PyCCR4-1::GFP) or to stage-specific cellular markers (CSP, ACP) are shown. Oocysts were imaged by live fluorescence. Scale bars are either 20 microns (oocysts), 5 microns (sporozoites), or 10 microns (asexual blood stage and liver stage parasites).

<u>S2 Figure (.PDF)</u>: (A) The four bioinformatically predictable CCR4 domain-containing proteins of *Plasmodium* species. The four proteins with identified exonuclease-endonuclease-phosphatase domains (shaded white rectangles) are shown to scale with their domain architecture, introns (gaps) and exons (rectangles). Also shown are E-values for their EEP domain based upon their alignment with CCR4 (PLN03144) via the Conserved Protein Domain Database. (B) The four bioinformatically predictable CCR4 domain-contains proteins from *P. yoelii*, the catalytically dead PyCCR4-1 (dPyCCR4-1), CCR4-1 from *P. falciparum*, and examples from *S. cerevisiae*, human, and mouse were aligned using EMBL Clustal Omega. Shown is the region around the catalytic residues of CCR4. Highlighted in red font are the two catalytic residues and in black highlighting with white font are the two residues that were changed to create the catalytic dead variant. (C-F) Genotyping PCR of (C) *pyccr4-1*^-^, (D) *pyccr4-2*^-^, (E) *pyccr4-3*^-^, and (F) *pyccr4-4*^-^ transgenic parasites. Successful genetic deletions were created using double homologous recombination of the targeting sequence consisting of ^~^750bp on either side of the ORF. Genotyping was performed by PCR on parasites cloned by limiting dilution using the primers indicated (listed in S7 Table). Independent clones were compared to a Py17XNL wild-type control, a no template control, and a plasmid positive control in parallel.

<u>S3 Figure (.PDF)</u>: (A) Asexual blood stage growth was monitored for two *pyccr4-1*^-^ transgenic clonal lines compared to a WT-GFP control line over the entire course of an infection. No significant difference in growth kinetics was observed. (B) Gametocyte counts were performed using flow cytometry. Asexual stage parasites were removed with two days of sulfadiazine treatment and WBC’s were removed using a cellulose column. PyDDD high and BIP + cells were scored as mature male gametocytes and DDD mid and BIP+ cells were scored as immature gametocytes and female gametocytes. No red blood cells were excluded in this analysis, and thus permitted measurement of gametocytemia. A PyDDD promoter driving GFP was used to establish gating of mature male gametocytes. PyDDD+ cells were FACS selected and observed to be male gametocytes (giemsa staining) that could undergo gametogenesis (exflagellation assay). (C) Genotyping PCR of dCCR4-1 transgenic parasites is shown. A successful replacement of the PyCCR4-1 catalytic residues were created using double homologous recombination to insert a C-terminal GFP tag and stop codon following the PyCCR4-1 stop codon. Genotyping was performed by PCR on parasites cloned by limiting dilution cloning using the primers indicated (listed in S7 Table). Independent clones were analyzed with a Py17XNL wild-type control, a no template control, and a plasmid positive control in parallel. Sequencing results are shown demonstrating the appropriate base change has occurred. This mutates the putative active residues to alanines. (D) Raw gametocytemia values are plotted. Error bars are standard error of the mean. (E) A mosquito feed was performed 2 days after the peak day of exflagellation (D7). The number of oocysts per infected mosquito on day seven post-infectious blood meal are plotted. Data represents at least 20 dissected mosquitoes per biological replicate conducted in triplicate. Error bars represent the standard error of the mean.

<u>S4 Figure (.PDF)</u>: (A) Genotyping PCR of *pycaf*^-^ transgenic parasites. An unsuccessful attempt at gene deletion by double homologous recombination using targeting sequences consisting of ^~^750bp on either side of the ORF is depicted. Genotyping was performed by PCR on parasites subjected to two rounds of drug selection using the primers indicated (listed in S7 Table). These parasites were compared to a Py17XNL wild-type control, a no template control, and a plasmid positive control in parallel. (B) A *P. falciparum* line carrying a *piggyBac* transposon inserted after the CAF1 deadenylase domain makes a truncated transcript. Schematic of RT-PCR primers aligned to the CAF1 ORF. The site of the *piggyBac* disruption is indicated by dotted line. RT-PCR results indicate wild-type NF54 and PfCAF1ΔC parasites both make transcript from the deadenylase domain (primer set P1; S7 Table). PfCAF1ΔC parasites do not make a full-length transcript (primer set P2; S7 Table). (C) Genotyping PCR of *pycaf1* disruptant transgenic parasites is shown. A successful disruption of *pycaf1* was created using double homologous recombination to insert a C-terminal GFP tag and stop codon following the CAF1 domain. Genotyping was performed by PCR on parasites cloned by limiting dilution cloning using the primers indicated (listed in S7 Table). Independent clones were analyzed with a Py17XNL wild-type control, a no template control, and a plasmid positive control in parallel.

<u>S5 Figure (.PDF)</u>: Expression of PyCAF1ΔC prevents its association with the CAF1/CCR4/NOT complex and its binding partners. A comparison of interactions by PyCCR4-1::GFP or PyCAF1ΔC1::GFP in blood stage schizonts was conducted to determine effects upon the assembly of the CAF1/CCR4/Not complex. (A) Using a very stringent SAINT score cutoff (0.1), only two proteins are in common between the PyCCR4-1::GFP and PyCAF1ΔC interactomes. Using an expanded SAINT score cutoff (0.35), relatively few interactions are still found to be in common. (B) Interaction of PyCCR4 with PyCAF1 and PyCAF1 with the NOT complex is abolished with PyCAF1 truncation (PyCAF1ΔC). Reported are proteins that interact with PyCCR4-1::GFP or PyCAF1ΔC::GFP using very stringent SAINT Scores (<0.1, unshaded), stringent SAINT scores (0.1 to 0.35, shaded light gray) and those with SAINT scores above 0.35 (shaded dark gray).

<u>S6 Figure (.PDF)</u>: (A) PyCAF1::GFP is diffusely expressed with some areas of higher intensity in asexual blood stages and in cytosolic granules in gametocytes. Representative images of asexual blood stages, and sexual stages treated with DAPI and antibodies to GFP (to detect PyCAF1::GFP) or to stage-specific cellular markers (ACP, alpha-tubulin, or human DDX6 that cross-reacts with DOZI) are shown. Scale bars are 5 microns. (B) PyCAF1ΔC::GFP is diffusely expressed in asexual blood stages and diffusely expressed with some areas of higher intensity in gametocytes. Representative images of asexual blood stages, and sexual stages treated with DAPI and antibodies to GFP (to detect PyCAF1::GFP) or to stage-specific cellular markers (ACP, alpha-tubulin, or human DDX6 that cross-reacts with DOZI) are shown. Scale bars are 5 microns.

<u>S7 Figure</u>: Extended data related to Figure 5. Control reactions lacking reverse transcriptase are provided in addition to the +RT experimental samples for all assays. Assessment of *gapdh* by cRT-PCR is also provided as a control.

<u>S8 Figure</u>: Sanger sequencing of cRT-PCR products from the circularized p28 transcript using primers that anneal within the coding sequence (upper case) permitted identification of the 5’ (red font, lower case) and 3’ UTRs (blue font, lower case), as well as the poly(A) tail (black font, lower case, underlined). Sequencing could not extend robustly through the poly(A) tail to provide an exact length from either forward or reverse sequencing primers (denoted by dashes). Sequences of the circularized gene product are provided from a cloned PCR product that is representative of UTRs from both wild-type and *pyccr4-1*^-^ samples.

<u>S1 Table (.XLSX)</u>: Measurements of transmission-related phenomena for each *pyccr4-1*^-^ clone is shown as averages for each replicate. Centers of movement/exflagellation centers are shown as an average of the number of exflagellating males per field in ten 400x fields. Prevalence of mosquito infections, oocyst counts, and sporozoite counts are all averages from at least 20 mosquitoes per replicate conducted in biological triplicate. Additional information on methods and data listed is provided in a README tab. Male and female gametocyte counts are shown for wild-type parasites expressing GFPmut2 from a disrupted *p230p* locus and *pyccr4-1*^-^ transgenic parasite lines. There is no appreciable difference in male gametocytemia between each line that would explain the extent of the decrease of male gametocytes that can activate into gametes on the peak day.

<u>S2 Table (.XLSX)</u>: The raw and annotated outputs from DEseq2 comparison of transcript abundance from four biological replicates of *pyccr4-1*^-^ parasites vs six biological replicates of Py17XNL wild-type parasites is provided. Additional information on data listed is provided in a README tab.

<u>S3 Table (.XLSX)</u>: Genes associated with translationally repressed transcripts in female gametocytes (31), male gamete-enriched proteins (32), and transcripts that are affected by deletion of *pyccr4-1* are listed. Comparisons of these lists were used to generate Venn diagrams (Figure 2). The Input tab is the gene ID’s from each input and the Output tab contains the independent and overlapping fields of the Venn diagram in Figure 2.

<u>S4 Table (.XLSX)</u>: The total proteomes of PY17XNL wild-type parasites and *pyccr4-1*^-^ parasites are provided as their RAW output, FDR 1% cutoff and a list of the *Plasmodium* proteins detected within the 1% FDR cutoff for each parasite type.

<u>S5 Table (.XLSX)</u>: Output files from TPP (The Trans-Proteomic Pipeline) are shown in individual tables for each control and experimental replicate. The output from the SAINT IP-MS data analysis is also shown with the PyCCR4-1::GFP bait protein manually added. Identified proteins are sorted by SAINT Scores, with highly stringent (< 0.1) being unshaded and stringent (0.1 to 0.35) hits shaded light grey, and all proteins with SAINT scores >0.35 shaded dark gray. The output files from TPP (The Trans-Proteomic Pipeline) are shown in individual tables for each control and experimental replicate. The output from the SAINT IP-MS data analysis is also shown with the PyCAF1ΔC::GFP bait protein manually added. Identified proteins are sorted by SAINT Scores, with highly stringent (< 0.1) being unshaded and stringent (0.1 to 0.35) hits shaded light grey, and all proteins with SAINT scores >0.35 shaded dark gray.

<u>S6 Table (.XLSX)</u>: Measurements of *P. falciparum* transmission-related phenotypes. Data are shown for each replicate. Additional information is provided in a README tab.

<u>S7 Table (.XLSX)</u>: Oligonucleotides that were used to generate and genotype transgenic parasites are shown. Upper case letters indicate nucleotides that match perfectly with the native genomic sequence, while lower case letters do not.

## REFERENCES

1. World Malaria Report: World Health Organization.; 2017. Available from: http://apps.who.int/iris/bitstream/handle/10665/259492/9789241565523-eng.pdf;jsessionid=70DDB5D8052E7F941829E6BAEC189ED0?sequence=1.

2. Smith RC, Vega-Rodriguez J, Jacobs-Lorena M. The Plasmodium bottleneck: malaria parasite losses in the mosquito vector. Mem Inst Oswaldo Cruz. 2014;109(5):644–61.

3. Rosenberg R, Wirtz RA, Schneider I, Burge R. An estimation of the number of malaria sporozoites ejected by a feeding mosquito. Trans R Soc Trop Med Hyg. 1990;84(2):209–12.

4. Munoz EE, Hart KJ, Walker MP, Kennedy MF, Shipley MM, Lindner SE. ALBA4 modulates its stage-specific interactions and specific mRNA fates during Plasmodium yoelii growth and transmission. Mol Microbiol. 2017.

5. Cui L, Lindner S, Miao J. Translational regulation during stage transitions in malaria parasites. Ann N Y Acad Sci. 2015;1342:1–9.

6. Lindner SE, Mikolajczak SA, Vaughan AM, Moon W, Joyce BR, Sullivan WJ, Jr., et al. Perturbations of Plasmodium Puf2 expression and RNA-seq of Puf2-deficient sporozoites reveal a critical role in maintaining RNA homeostasis and parasite transmissibility. Cellular microbiology. 2013;15(7):1266–83.

7. Muller K, Matuschewski K, Silvie O. The Puf-family RNA-binding protein Puf2 controls sporozoite conversion to liver stages in the malaria parasite. PLoS One. 2011;6(5):e19860.

8. Gomes-Santos CS, Braks J, Prudencio M, Carret C, Gomes AR, Pain A, et al. Transition of Plasmodium sporozoites into liver stage-like forms is regulated by the RNA binding protein Pumilio. PLoS Pathog. 2011;7(5):e1002046.

9. Painter HJ, Campbell TL, Llinas M. The Apicomplexan AP2 family: integral factors regulating Plasmodium development. Mol Biochem Parasitol. 2011;176(1):1–7.

10. Painter HJ, Carrasquilla M, Llinas M. Capturing in vivo RNA transcriptional dynamics from the malaria parasite Plasmodium falciparum. Genome Res. 2017;27(6):1074–86.

11. Guerreiro A, Deligianni E, Santos JM, Silva PA, Louis C, Pain A, et al. Genome-wide RIP-Chip analysis of translational repressor-bound mRNAs in the Plasmodium gametocyte. Genome Biol. 2014;15(11):493.

12. Heather J Painter NCC, Aswathy Sebastian, Istvan Albert, John D Storey, Manuel Llinás. Real-time in vivo Global Transcriptional Dynamics During Plasmodium falciparum Blood-stage Development bioRxiv2018 [Available from: https://www.biorxiv.org/content/early/2018/02/14/265975.

13. Mair GR, Braks JA, Garver LS, Wiegant JC, Hall N, Dirks RW, et al. Regulation of sexual development of Plasmodium by translational repression. Science. 2006;313(5787):667–9.

14. Mair GR, Lasonder E, Garver LS, Franke-Fayard BM, Carret CK, Wiegant JC, et al. Universal features of post-transcriptional gene regulation are critical for Plasmodium zygote development. PLoS Pathog. 2010;6(2):e1000767.

15. Paton MG, Barker GC, Matsuoka H, Ramesar J, Janse CJ, Waters AP, et al. Structure and expression of a post-transcriptionally regulated malaria gene encoding a surface protein from the sexual stages of Plasmodium berghei. Mol Biochem Parasitol. 1993;59(2):263–75.

16. Parker R, Sheth U. P bodies and the control of mRNA translation and degradation. Mol Cell. 2007;25(5):635–46.

17. Lindner SE, Swearingen KE, Harupa A, Vaughan AM, Sinnis P, Moritz RL, et al. Total and putative surface proteomics of malaria parasite salivary gland sporozoites. Molecular & cellular proteomics : MCP. 2013;12(5):1127–43.

18. Balagopal V, Parker R. Polysomes, P bodies and stress granules: states and fates of eukaryotic mRNAs. Curr Opin Cell Biol. 2009;21(3):403–8.

19. Collart MA. The Ccr4-Not complex is a key regulator of eukaryotic gene expression. Wiley Interdiscip Rev RNA. 2016;7(4):438–54.

20. Miller JE, Reese JC. Ccr4-Not complex: the control freak of eukaryotic cells. Crit Rev Biochem Mol Biol. 2012;47(4):315–33.

21. Niinuma S, Fukaya T, Tomari Y. CCR4 and CAF1 deadenylases have an intrinsic activity to remove the post-poly(A) sequence. RNA. 2016;22(10):1550–9.

22. Collart MA, Panasenko OO. The Ccr4--not complex. Gene. 2012;492(1):42–53.

23. Ukleja M, Cuellar J, Siwaszek A, Kasprzak JM, Czarnocki-Cieciura M, Bujnicki JM, et al. The architecture of the Schizosaccharomyces pombe CCR4-NOT complex. Nature communications. 2016;7:10433.

24. Balu B, Maher SP, Pance A, Chauhan C, Naumov AV, Andrews RM, et al. CCR4-associated factor 1 coordinates the expression of Plasmodium falciparum egress and invasion proteins. Eukaryotic cell. 2011;10(9):1257–63.

25. Balu B, Singh N, Maher SP, Adams JH. A genetic screen for attenuated growth identifies genes crucial for intraerythrocytic development of Plasmodium falciparum. PLoS One. 2010;5(10):e13282.

26. Bushell E, Gomes AR, Sanderson T, Anar B, Girling G, Herd C, et al. Functional Profiling of a Plasmodium Genome Reveals an Abundance of Essential Genes. Cell. 2017;170(2):260–72 e8.

27. Reddy BP, Shrestha S, Hart KJ, Liang X, Kemirembe K, Cui L, et al. A bioinformatic survey of RNA-binding proteins in Plasmodium. BMC Genomics. 2015;16(1):890.

28. Cooke A, Prigge A, Wickens M. Translational repression by deadenylases. J Biol Chem. 2010;285(37):28506–13.

29. Guntur AR, Kawai M, Le P, Bouxsein ML, Bornstein S, Green CB, et al. An essential role for the circadian-regulated gene nocturnin in osteogenesis: the importance of local timekeeping in skeletal homeostasis. Ann N Y Acad Sci. 2011;1237:58–63.

30. Temme C, Zhang L, Kremmer E, Ihling C, Chartier A, Sinz A, et al. Subunits of the Drosophila CCR4-NOT complex and their roles in mRNA deadenylation. RNA. 2010;16(7):1356–70.

31. Jain S, Wheeler JR, Walters RW, Agrawal A, Barsic A, Parker R. ATPase-Modulated Stress Granules Contain a Diverse Proteome and Substructure. Cell. 2016;164(3):487–98.

32. Guo L, Kim HJ, Wang H, Monaghan J, Freyermuth F, Sung JC, et al. Nuclear-Import Receptors Reverse Aberrant Phase Transitions of RNA-Binding Proteins with Prion-like Domains. Cell. 2018;173(3):677–92 e20.

33. Sievers F, Higgins DG. Clustal omega. Curr Protoc Bioinformatics. 2014;48:3 13 1–6.

34. Dupressoir A, Morel AP, Barbot W, Loireau MP, Corbo L, Heidmann T. Identification of four families of yCCR4-and Mg2+-dependent endonuclease-related proteins in higher eukaryotes, and characterization of orthologs of yCCR4 with a conserved leucine-rich repeat essential for hCAF1/hPOP2 binding. BMC Genomics. 2001;2:9.

35. Raisch T, Bhandari D, Sabath K, Helms S, Valkov E, Weichenrieder O, et al. Distinct modes of recruitment of the CCR4-NOT complex by Drosophila and vertebrate Nanos. EMBO J. 2016;35(9):974–90.

36. Sandler H, Kreth J, Timmers HT, Stoecklin G. Not1 mediates recruitment of the deadenylase Caf1 to mRNAs targeted for degradation by tristetraprolin. Nucleic Acids Res. 2011;39(10):4373–86.

37. Stowell JA, Webster MW, Kogel A, Wolf J, Shelley KL, Passmore LA. Reconstitution of Targeted Deadenylation by the Ccr4-Not Complex and the YTH Domain Protein Mmi1. Cell Rep. 2016;17(8):1978–89.

38. Schwach F, Bushell E, Gomes AR, Anar B, Girling G, Herd C, et al. PlasmoGEM, a database supporting a community resource for large-scale experimental genetics in malaria parasites. Nucleic Acids Res. 2015;43(Database issue):D1176–82.

39. Anders S, Huber W. Differential expression analysis for sequence count data. Genome Biol. 2010;11(10):R106.

40. Noble WS. How does multiple testing correction work? Nat Biotechnol. 2009;27(12):1135–7.

41. Ikadai H, Shaw Saliba K, Kanzok SM, McLean KJ, Tanaka TQ, Cao J, et al. Transposon mutagenesis identifies genes essential for Plasmodium falciparum gametocytogenesis. Proc Natl Acad Sci U S A. 2013;110(18):E1676–84.

42. Farber V, Erben E, Sharma S, Stoecklin G, Clayton C. Trypanosome CNOT10 is essential for the integrity of the NOT deadenylase complex and for degradation of many mRNAs. Nucleic Acids Res. 2013;41(2):1211–22.

43. Yeoh LM, Goodman CD, Mollard V, McFadden GI, Ralph SA. Comparative transcriptomics of female and male gametocytes in Plasmodium berghei and the evolution of sex in alveolates. BMC Genomics. 2017;18(1):734.

44. Lasonder E, Rijpma SR, van Schaijk BC, Hoeijmakers WA, Kensche PR, Gresnigt MS, et al. Integrated transcriptomic and proteomic analyses of P. falciparum gametocytes: molecular insight into sex-specific processes and translational repression. Nucleic Acids Res. 2016;44(13):6087–101.

45. Otto TD, Bohme U, Jackson AP, Hunt M, Franke-Fayard B, Hoeijmakers WA, et al. A comprehensive evaluation of rodent malaria parasite genomes and gene expression. BMC Biol. 2014;12:86.

46. Lopez-Barragan MJ, Lemieux J, Quinones M, Williamson KC, Molina-Cruz A, Cui K, et al. Directional gene expression and antisense transcripts in sexual and asexual stages of Plasmodium falciparum. BMC Genomics. 2011;12:587.

47. Young JA, Fivelman QL, Blair PL, de la Vega P, Le Roch KG, Zhou Y, et al. The Plasmodium falciparum sexual development transcriptome: a microarray analysis using ontology-based pattern identification. Mol Biochem Parasitol. 2005;143(1):67–79.

48. Bunnik EM, Batugedara G, Saraf A, Prudhomme J, Florens L, Le Roch KG. The mRNA-bound proteome of the human malaria parasite Plasmodium falciparum. Genome Biol. 2016;17(1):147.

49. Halter D, Collart MA, Panasenko OO. The Not4 E3 ligase and CCR4 deadenylase play distinct roles in protein quality control. PLoS One. 2014;9(1):e86218.

50. Duy DL, Suda Y, Irie K. Cytoplasmic deadenylase Ccr4 is required for translational repression of LRG1 mRNA in the stationary phase. PLoS One. 2017;12(2):e0172476.

51. Woolstencroft RN, Beilharz TH, Cook MA, Preiss T, Durocher D, Tyers M. Ccr4 contributes to tolerance of replication stress through control of CRT1 mRNA poly(A) tail length. J Cell Sci. 2006;119(Pt 24):5178–92.

52. Eulalio A, Huntzinger E, Nishihara T, Rehwinkel J, Fauser M, Izaurralde E. Deadenylation is a widespread effect of miRNA regulation. RNA. 2009;15(1):21–32.

53. Methods in Malaria Research 2013. Available from: https://www.beiresources.org/portals/27MR4/MethodsInMalariaResearch-6thedition.pdf.

54. Carter R. The Culture and Preparation of Gametocytes of Plasmodium falciparum for Immunochemical, Molecular, and Mosquito Infectivity Studies. In: Hyde JE, editor. Protocols in Molecular Parasitology. Methods in Molecular Biology1993. p. 67–88.

55. Miller JL, Harupa A, Kappe SH, Mikolajczak SA. Plasmodium yoelii macrophage migration inhibitory factor is necessary for efficient liver-stage development. Infect Immun. 2012;80(4):1399–407.

56. Reece SE, Ali E, Schneider P, Babiker HA. Stress, drugs and the evolution of reproductive restraint in malaria parasites. Proc Biol Sci. 2010;277(1697):3123–9.

57. Mikolajczak SA, Vaughan AM, Kangwanrangsan N, Roobsoong W, Fishbaugher M, Yimamnuaychok N, et al. Plasmodium vivax liver stage development and hypnozoite persistence in human liver-chimeric mice. Cell Host Microbe. 2015;17(4):526–35.

58. Deutsch EW, Mendoza L, Shteynberg D, Slagel J, Sun Z, Moritz RL. Trans-Proteomic Pipeline, a standardized data processing pipeline for large-scale reproducible proteomics informatics. Proteomics Clin Appl. 2015;9(7-8):745–54.

59. Mellacheruvu D, Wright Z, Couzens AL, Lambert JP, St-Denis NA, Li T, et al. The CRAPome: a contaminant repository for affinity purification-mass spectrometry data. Nat Methods. 2013;10(8):730–6.

60. Shteynberg D, Deutsch EW, Lam H, Eng JK, Sun Z, Tasman N, et al. iProphet: multi-level integrative analysis of shotgun proteomic data improves peptide and protein identification rates and error estimates. Mol Cell Proteomics. 2011;10(12):M111 007690.

61. Choi H, Larsen B, Lin ZY, Breitkreutz A, Mellacheruvu D, Fermin D, et al. SAINT: probabilistic scoring of affinity purification-mass spectrometry data. Nat Methods. 2011;8(1):70–3.

62. Anders S, Pyl PT, Huber W. HTSeq--a Python framework to work with high-throughput sequencing data. Bioinformatics. 2015;31(2):166–9.

63. Love MI, Huber W, Anders S. Moderated estimation of fold change and dispersion for RNA-seq data with DESeq2. Genome Biol. 2014;15(12):550.

64. Team RC. R: A language and environment for statistical computing.: R Foundation for Statistical Computing, Vienna, Austria. ; 2016.

65. Vizcaino JA, Csordas A, del-Toro N, Dianes JA, Griss J, Lavidas I, et al. 2016 update of the PRIDE database and its related tools. Nucleic Acids Res. 2016;44(D1):D447–56.

66. Talman AM, Prieto JH, Marques S, Ubaida-Mohien C, Lawniczak M, Wass MN, et al. Proteomic analysis of the Plasmodium male gamete reveals the key role for glycolysis in flagellar motility. Malar J. 2014;13:315.

67. Bai Y, Salvadore C, Chiang YC, Collart MA, Liu HY, Denis CL. The CCR4 and CAF1 proteins of the CCR4-NOT complex are physically and functionally separated from NOT2, NOT4, and NOT5. Mol Cell Biol. 1999;19(10):6642–51.

68. Bhaskar V, Basquin J, Conti E. Architecture of the ubiquitylation module of the yeast Ccr4-Not complex. Structure. 2015;23(5):921–8.

69. Petit AP, Wohlbold L, Bawankar P, Huntzinger E, Schmidt S, Izaurralde E, et al. The structural basis for the interaction between the CAF1 nuclease and the NOT1 scaffold of the human CCR4-NOT deadenylase complex. Nucleic Acids Res. 2012;40(21):11058–72.

70. Kelley LA, Mezulis S, Yates CM, Wass MN, Sternberg MJ. The Phyre2 web portal for protein modeling, prediction and analysis. Nat Protoc. 2015;10(6):845–58.

71. States DJ, Gish W. Combined use of sequence similarity and codon bias for coding region identification. J Comput Biol. 1994;1(1):39–50.

